# Interfamily co-transfer of sensor and helper NLRs extends immune receptor functionality between angiosperms

**DOI:** 10.1101/2024.12.11.627876

**Authors:** XiaoFei Du, Maheen Alam, Kamil Witek, Lesley Milnes, James Houghton, Xiao Lin, Yu Zhang, Fuhao Cui, Wenxian Sun, Jonathan D.G. Jones, Hailong Guo

## Abstract

Plant nucleotide-binding, leucine-rich repeat (NLR) immune receptors recognize pathogen effectors and activate defense. NLR genes can be non-functional in distantly related plants (restricted taxonomic functionality, RTF). Here, we enable Solanaceae NLR gene function in rice, soybean and Arabidopsis by co-delivering sensor NLR genes with their cognate NRC-type helper NLRs. In soybean protoplasts and in Arabidopsis plants, *Solanum americanum* Rpi-amr1, Rpi-amr3 and pepper Bs2 sensor NLRs confer cognate effector responsiveness if co-expressed with NRC helper NLRs. Rice carrying pepper Bs2 and NRCs recognizes the conserved effector, AvrBs2, and resists an important pathogen, *Xanthomonas oryzae* pv. *oryzicola* for which no resistance gene is available in rice. Rice lines carrying sensor and helper NLR genes otherwise resemble wild-type, with unaltered basal resistance or field fitness. Thus, interfamily co-transfer of sensor and helper NLRs can broaden utility of sensor NLRs, extending the tools available to control diseases of rice, soybean, Brassicas and other crops.

## Introduction

Plant pathogens cause devastating diseases in plants and pose a serious threat to global crop production (Savary et al., 2019). Plant defenses against microbial infections are initiated upon perception of pathogen-derived molecules by cell surface-localized pattern-recognition receptors (PRRs) and intracellular nucleotide-binding leucine-rich repeat receptors (NLRs) (Ngou et al., 2022). PRRs detect apoplastic microbe- or pathogen-associated molecular pattern (MAMP/PAMP) molecules, or host-derived damage-associated molecular patterns (DAMPs), and mount pattern-triggered immunity (PTI) (Boutrot and Zipfel, 2017). Adapted pathogens secrete effector proteins into host cells that promote virulence, enabling pathogen proliferation. Plants have evolved intracellular nucleotide-binding and leucine-rich repeat (NLR) immune receptors that detect effectors (either directly, or indirectly by monitoring the effector-mediated modification of host proteins) and activate effector-triggered immunity (ETI) (Jones et al., 2016). ETI often culminates in rapid host cell death localized to the infection sites, termed the Hypersensitive Response (HR) that contributes to pathogen growth restriction.

Most NLR immune receptors share a conserved tripartite domain architecture: an N-terminal signaling domain, a central nucleotide-binding (NB-ARC) domain and a C-terminal leucine-rich repeat (LRR) domain. The N-terminal domains of NLRs usually comprise either (i) N-terminal Toll/interleukin-1 receptor/R protein (TIR) domain (TNLs) (ii) coiled-coil (CC) domain (CNLs), or (iii) Resistance to Powdery mildew8 (RPW8)-like domains (RNLs). Recent breakthroughs reveal mechanisms of recognition, activation and signaling of plant NLRs (Contreras et al., 2023a; Wang and Chai, 2020). NLR proteins oligomerize upon effector recognition, imposing induced proximity on their N-terminal signaling domains, activating downstream immune signaling (Forderer et al., 2022; Ma et al., 2020; Martin et al., 2020; Wang et al., 2019a; Wang et al., 2019b). Since NLRs from one plant species can recognize effectors in pathogens of other species, NLR transfer between species might be useful to elevate crop resistance (Tai et al 1999).

Understanding plant immune receptors can facilitate enhancing disease resistance in crops (Kawashima et al., 2016; Liu et al., 2021; Luo et al., 2021). The Arabidopsis PRR EFR recognizes bacterial translation elongation factor EF-Tu and elevates resistance to *Ralstonia solanacearum* when transferred to Solanaceae (Lacombe et al., 2010). However, interfamily transfer of NLR genes sometimes only confers resistance in closely related plant species (Ghislain et al., 2019; Hatta et al., 2021; Kawashima et al., 2016; Witek et al., 2016). This phenomenon has been called restricted taxonomic functionality (RTF) (Tai et al., 1999). An emerging paradigm in plant immunity is that many sensor NLRs require a partner NLR or “helper NLR(s)” (Adachi et al., 2019b; Cesari et al., 2014). Thus, RTF for a sensor NLR could result from a requirement for the appropriate helper/partner NLR (Lapin et al., 2019). In Solanaceae, over half the NLRs are phylogenetically related and belong to the NRC (NLR required for cell death) superclade (Wu et al., 2017). NRC helper NLRs are only found in asterids and are absent from rosids such as Arabidopsis and soybean and from monocots such as rice (Sakai et al., 2024).

The RTF constraint was first discovered when pepper Bs2, that recognizes the *Xanthomonas campestris* pv. *vesicatoria* (*Xcv*) effector AvrBs2, was shown to function in closely related tomato but not in distantly related Arabidopsis (Tai et al., 1999). The AvrBs2 effector is widely distributed in various *Xanthomonas* pathovars that infect rosid crops such as cassava and Brassicas and monocot crops such as rice (Li et al., 2015; Medina et al., 2018). Similarly, the NRC-dependent *Rpi-amr3* (*Resistance to Phytophthora infestans Solanum americanum 3*) recognizes a *Phytophthora infestans* effector (AvrAmr3) that is found in many *Phytophthora* pathogen species infecting rosid crops such soybean, Brassicas and cacao (Lin et al., 2022; Oh et al., 2024). Extending recognition of these pathogens via *Bs2* or *Rpi-amr3* into these rosid crops could enhance crop disease resistance.

We investigated if co-delivery of sensor and NRC helper NLRs can overcome RTF. We found *Bs2, Rpi-amr1* and *Rpi-amr3* function in transient assays in soybean and in stable Arabidopsis transgenic lines if NRCs are co-provided. We used this approach to address a major rice (*Oryza sativa*) disease. Bacterial leaf streak (BLS) caused by *Xanthomonas oryzae* pv. *oryzicola* (*Xoc*) is a serious threat to rice production worldwide. *Xoc* enters through stomata or wounds, colonizes the mesophyll apoplast and reduces yield. However, no endogenous BLS resistance genes have been identified in rice. AvrBs2 is highly conserved in many *Xanthomonas* pathovars and required for the virulence of *Xoc*, *Xcv* and *Xanthomonas axonopodis* pv. *manihotis* (*Xam*) (Li et al., 2015; Medina et al., 2018; Zhao et al., 2011). We tested whether transfer of pepper *Bs2*, which also recognizes AvrBs2 allele from *Xoc*, into rice would confer BLS resistance and found NRC-dependent efficacy, without affecting basal immunity or affecting rice growth and yield. These data demonstrate for the first time that co-transfer of sensor and helper NLRs breaks RTF and enable new paths to elevate resistance against many pathogens of rosid and monocot crops through redeploying known NLR genes.

## Results

### Helper NLR-dependent cell death activation by Solanaceae sensor NLRs in rosid and rice transient assays

AvrBs2 in *Xcv* and *Xoc* possess high sequence identity and functional similarity (Li et al., 2015). To investigate whether orthologous AvrBs2*^Xoc^*is recognized by Bs2, we cloned AvrBs2*^Xoc^* into an expression vector with the 35S promoter and performed transient expression assays in *N. benthamiana*. Like AvrBs2*^Xcv^*, AvrBs2*^Xoc^* alone does not trigger HR in *N. benthamiana*, but AvrBs2*^Xoc^*can induce HR when co-expressed with Bs2 (Figure 1A). These results were confirmed by electrolyte leakage induced by co-expression of AvrBs2*^Xoc^* with Bs2 but not with GFP control (Figure 1B), suggesting that AvrBs2*^Xoc^* is also recognized by Bs2.

**Figure 1.**
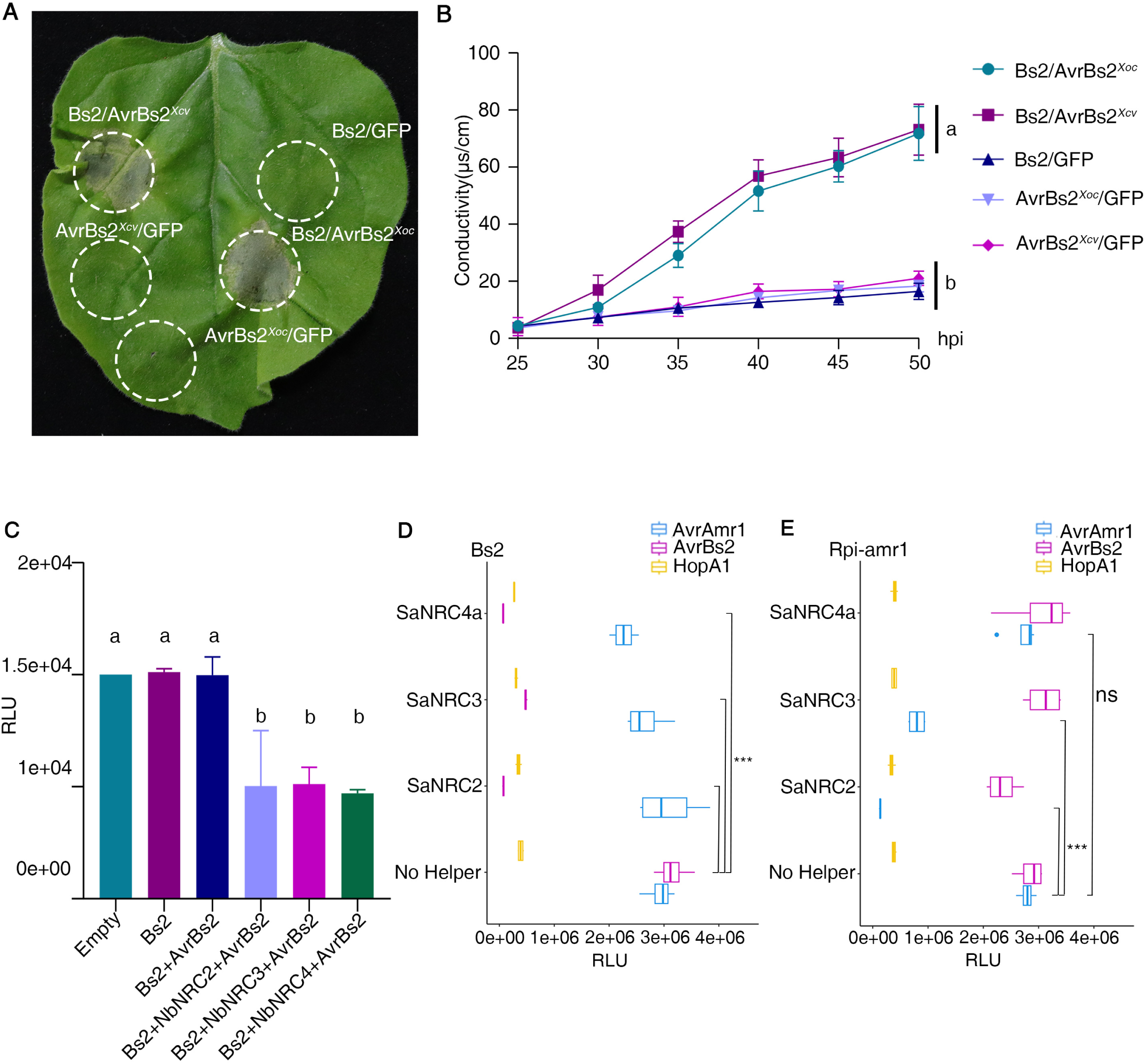
Transient assays verifying sufficiency of NRC helpers to support sensor NLR function in non-asterids. **(A)** HR induction by expressing Bs2 in the presence of AvrBs2*^Xoc^* or AvrBs2*^Xcv^*. The *N. benthamiana (Nb)* leaves were infiltrated with *Agrobacterium* strain GV3101 carrying the indicated constructs and the HR phenotype was photographed at 5 days post-infiltration (dpi). Three biological replicates were performed with similar results. **(B)** Electrolyte leakage in *Nb* leaves was monitored from 20 to 50 hours post infiltration (hpi). Upon expression of the same constructs combinations as indicated in (A) by agroinfiltration, electrolyte leakage was measured every five hours from 20 to 50 hpi. Error bars were calculated from three replicates per time point and per construct. Letters (a–b) represent groups with significant differences [p<0.05, Tukey’s honest significant difference (HSD) test]. **(C)** Luciferase activity assay measuring cell death in rice protoplasts upon coexpression of AvrBs2 and Bs2 in the presence or absence of NRCs. The indicated constructs were co-transfected with a luciferase reporter vector, and luciferase activity was measured 24h post transfection. This experiment was independently repeated three times with similar results. (**D-E**) Co-expression of Bs2 (D) or Rpi-amr1 (E) with AvrAmr1, AvrBs2 or cell death control HopA1, in the presence or absence of NRCs, in soybean protoplasts. The protoplasts were also transfected with Luciferase reporter. Luciferase activity was measured in relative light units 48 h post transfection. The experiments were repeated independently three times with similar results.

We then tested whether Bs2 confers AvrBs2*^Xoc^* recognition in rice using protoplast transient expression luciferase eclipse assays. No reduction of luciferase activity was observed upon co-transfection of 35S:LUC with Bs2 and AvrBs2*^Xoc^* (Figure 1C). Bs2 function requires the *N. benthamiana* helper NLRs NbNRC2, NbNRC3, or NbNRC4 that are absent from monocot genomes (Wu et al., 2017). We tested if co-delivery of Bs2 and helper NLRs could enable immune response to AvrBs2*^Xoc^*. Transient codelivery of Bs2 with AvrBs2*^Xoc^* significantly reduced LUC reporter activity in the presence of NbNRC2, NbNRC3, or NbNRC4 (Figure 1C), consistent with transient expression assays in *N. benthamiana*. Transient overexpression of AvrBs2*^Xoc^* with NRCs in protoplasts resulted in no reduction in luciferase activity (Figure S1A). Collectively, transient assays in *N. benthamiana* and rice protoplasts indicate that AvrBs2*^Xoc^*is recognized by Bs2 in a NRC helpers-dependent manner.

To test if a similar approach can relieve RTF with other sensor and helper NLRs and in rosid plants, similar experiments were conducted in soybean protoplasts. Either Bs2, or the *Solanum americanum* late blight resistance gene *Rpi-amr1* whose function is dependent on NRC2 or NRC3 but not NRC4 (Witek et al., 2021), were co-delivered with 35S:LUC and with either *S. americanum* SaNRC2, SaNRC3 or SaNRC4a in the presence or absence of recognized effector. The recognized effector HopA1 was used as a positive control. Luciferase signal is represented on a log scale as Relative Light Units (RLU). When Bs2 was co-delivered with SaNRCs, RLU signal was suppressed in the presence of AvrBs2 but not AvrAmr1 (Lin et al., 2020), and vice versa when Rpi-amr1 was co-delivered with SaNRC2, SaNRC3 but not with SaNRC4a (Fig 1D-E). These verify the findings of Fig 1A, B, C for an additional angiosperm clade and for additional sensor and helper NLRs. Another sensor NLR, Rpi-amr3 whose resistance is dependent on NRC2, NRC3 or NRC4 (Lin et al., 2022; Witek et al., 2016), also showed SaNRC2/3/4-dependent AvrAmr3 recognition and luciferase eclipse (Fig S1B).

Additionally Rpi-amr3+SaNRC2 were co-expressed with 35S:LUC and with either AvrAmr1 or AvrAmr3 in *Arabidopsis thaliana* protoplasts. When Rpi-amr3 was co-delivered with SaNRC2 and AvrAmr3 the RLU signal was suppressed, whereas no signal reduction was observed in the presence of AvrAmr1 (Fig S1C). Collectively, these data indicate that for Rpi-amr1, Rpi-amr3 and Bs2, sensor-helper stacks confer recognition of their cognate effector in non-Asterid plant-derived protoplasts.

### Stable Arabidopsis lines carrying sensor and helper NLRs show effector-dependent HR

To test functionality of Solanaceae NLRs in non-Asterid *Arabidopsis thaliana* plants, we generated constructs that either carry sensor NLR or sensor+helper NLR constructs (Fig S2A). We assembled pUbi10:Rpi-amr3-FLAG and pUbi10:Bs2-FLAG with 35S:NbNRC2-MYC and also assembled pUbi10:Rpi-amr3-FLAG with 35S:SaNRC2-MYC (Fig S2A).The functionality and expression of these constructs were confirmed in transient assays in *Nb nrc2/3/4* CRISPR mutant plants (Fig S2B-C). Bs2 or Rpi-amr3 alone did not support effector-dependent cell death in the absence of an NRC (Fig S2B) but co-delivery of Bs2 and NbNRC2 or Rpi-amr3 and NbNRC2 constructs with AvrBs2 or AvrAmr3 respectively resulted in cell death.

These constructs were subsequently transformed into Arabidopsis Col-0 plants and for each construct, two or more independent lines were produced and tested. To assess if the sensor together with helper NLR constructs function in stable transgenic Arabidopsis, we tested independent lines by delivering the cognate effectors using the effector detector vector (EDV) (Sohn et al., 2007). AvrBs2 (aa 62-714) and AvrAmr3 (lacking signal peptide and RXLR motif, aa59-339) were cloned in pEDV and transformed in *P. syringae* pv. *tomato* DC3000, D36E (Fig S2D) (Wei et al., 2018). EDV efficacy was tested in *N. benthamiana* (Fig S2E). Two days after agroinfiltration of either sensor NLR or empty vector, leaves were infiltrated with D36E carrying the pEDV construct. Cell death was visible in the infiltrated area where the cognate sensor NLR had been infiltrated; we concluded that these effectors can be delivered via T3SS (Fig S2E).

Independent transgenic lines of Arabidopsis expressing sensor together with helper NLRs or sensor NLRs alone (Fig S2F) were tested for effector recognition. Figure 2 shows that lines co-transformed with *Rpi-amr3* and *NbNRC2* or *Rpi-amr3* and *SaNRC2* recognize AvrAmr3 delivered from DC3000 D36E; responsiveness culminates in cell death. No cell death was observed in lines transformed with *Rpi-amr3* alone. D36E carrying AvrRpt2 was used as a positive control and D36E carrying empty vector as a negative control. However, AvrBs2 recognition in lines carrying Bs2-NbNRC2 resulted in a weaker HR (Fig S2G-I). We speculate that the difference between transient assay and stable line phenotypes might be due to differences in NRC paralog efficacy for different sensors. We conclude that the effector is recognized in transgenic plants carrying the Solanaceae sensor together with helper NLR construct but not in sensor NLR alone lines.

**Figure 2.**
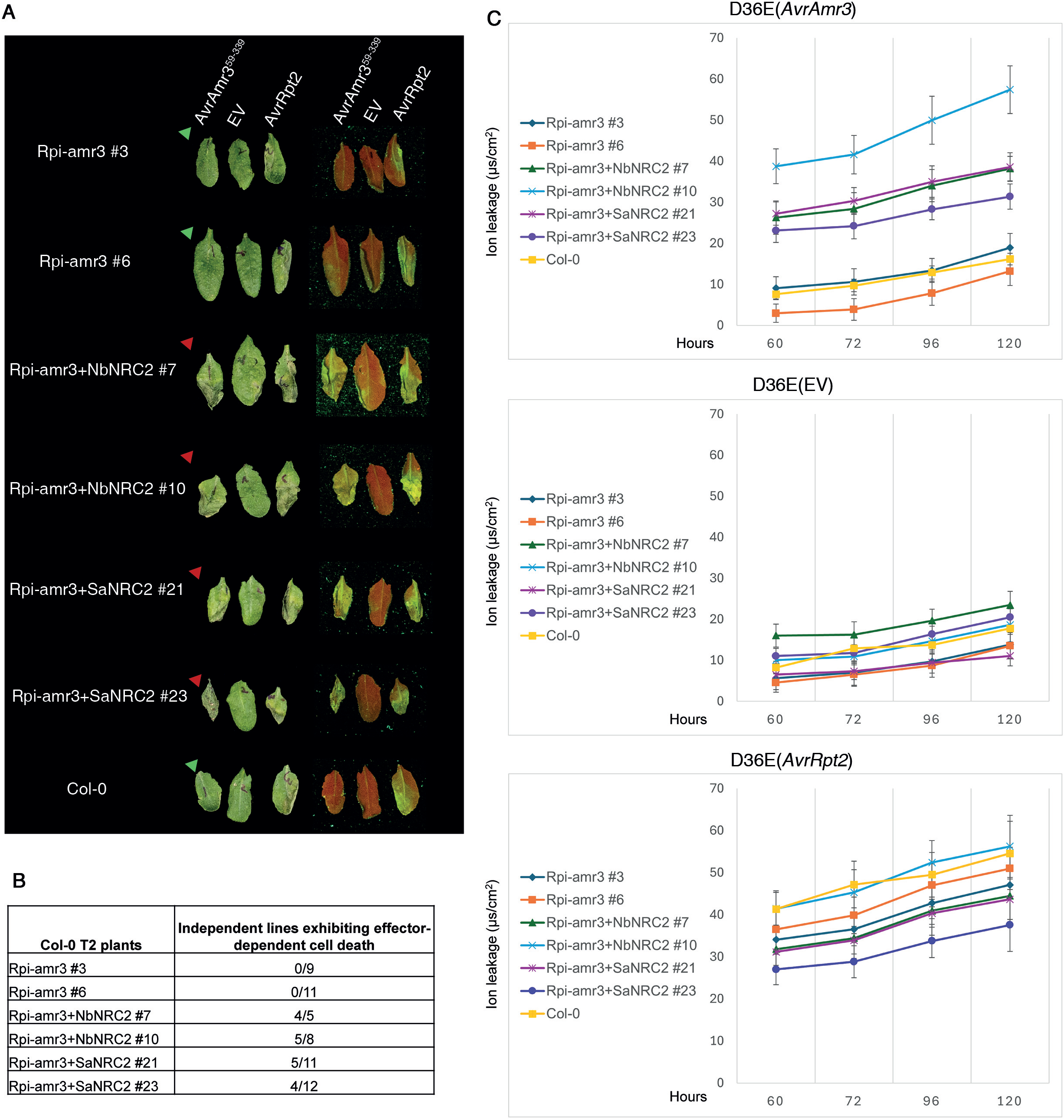
Effector recognition by Solanaceae Rpi-amr3/NRC2 sensor-helper pair results in cell death in *A. thaliana*. **(A)** Independent T2 lines were infiltrated with *P. syringae* D36E strains that deliver by T3SS either AvrAmr3, empty vector or AvrRpt2. Effector recognition by the Rpi-amr3/NRC2 sensor-helper pair resulted in cell death (indicated by red arrows-HR); whereas no cell death was observed in the sensor alone T2 plants (indicated by green arrows-no HR). Images were taken 60 hours post-inoculation (hpi). Panel on the right shows images of the same leaves taken under UV light. Cell death can be observed in green in the infiltrated area, while in the absence of cell death the infiltrated area is red. **(B)** Occurrence of cell death upon effector recognition in independent T2 lines of Col-0 that express Rpi-amr3+NRC2. **(C)** Cell death in leaves was quantified by measuring electrolyte leakage. Data shown were collected from 2 independent experiments with at least 3 technical replicates per transgenic line. Error bars represent standard error of the mean (SEM)

### Bs2 confers resistance in transgenic rice against *Xoc* carrying AvrBs2 in a NRC2, NRC3, or NRC4 dependent manner

Extending functionality of Solanaceae NLRs to non-Asterid plants provides a strategy to broaden the use of lineage-specific NLR genes across different plant families for disease control. *Xoc*, causes the destructive rice bacterial disease BLS. However, no endogenous BLS resistance gene has been found in rice. The recognition of AvrBs2*^Xoc^* by Bs2 in the presence of NbNRCs led us to investigate whether transgenic rice carrying Bs2 and NRCs were resistant to *Xoc*. We generated transgenic rice expressing Bs2 with either NbNRC2, NbNRC3, or NbNRC4 as well as a combination of the NRC-helpers NbNRC2/3/4. We employed Golden Gate cloning to stack NLR genes and chose synthetic C-terminal fusion epitope tags FLAG, HA, MYC and V5 for detecting Bs2, NbNRC2, NbNRC3 and NbNRC4 protein expression, respectively (Figure S3A-C). We introduced these constructs individually into ZH11 rice cultivar by *Agrobacterium*-mediated transformation. For each transgene cassette, multiple independent transgenic rice lines were generated and immunoblot analysis confirmed that proteins of interest were correctly expressed in the transgenic rice lines except for NbNRC3, which was not detected in the immunoblots but was only detectable after immunoprecipitation (Figure S3A-C). This result aligns with previously published observations that NbNRC3 doesn’t express well even in *N. benthamiana* compared with NbNRC2 and NbNRC4 (Derevnina et al., 2021).

To evaluate BLS resistance of transgenic rice lines, *Xoc* strain RS105 was inoculated into their leaves and wild-type ZH11. Leaves of wild-type ZH11 exhibited characteristic water-soaking lesions expanding from the inoculation sites, suggesting that *Xoc* is fully virulent on ZH11 (Figure 3A-B). Transgenic rice plants co-expressing *Bs2* and *NbNRC3* or *NbNRC4* were fully resistant against *Xoc* as no or only confined lesions were observed after infection (Figure 3A-B), suggesting that *Xoc* became avirulent on these transgenic rice plants. However, the *Bs2* and *NbNRC2* co-expressing lines exhibited partial/quantitative resistance against *Xoc* compared with transgenic lines co-expressing *Bs2* with either *NbNRC3* or *NbNRC4* (Figure 3A-B). The partial resistance in the *Bs2* and *NbNRC2* co-expressing lines is not a consequence of reduced Bs2 and NRC2 protein accumulation because Bs2 could be detected in all of these lines at similar levels and NbNRC2 accumulated even higher compared to NbNRC3 (Figure S3A-C). These results together suggest that *NbNRC* genes show different efficacy when heterologously expressed. Consistent with the reduced lesion lengths, growth of *Xoc* was markedly decreased in these transgenic lines, compared to the wild-type ZH11 (Figure 3C). In parallel, we generated transgenic rice co-expressing *Bs2* and the combination of the NRC-helpers *NbNRC2/3/4*(Figure S3D). Inoculation assays with *Xoc* demonstrated that these lines exhibited a similar level of resistance against *Xoc* as the transgenic lines co-expressing *Bs2* with *NRC3* or *NRC4* individually without a synergistic effect (Figure 3A-C), in contrast to the additive effects for stacks of multiple sensor NLRs (Ghislain et al., 2019; Xiao et al., 2016). Δ*AvrBs2* mutant *Xoc* strain in which the *AvrBs2* gene has been disrupted was still able to cause disease on all rice lines tested although Δ*AvrBs2* mutant *Xoc* strain is less virulent on rice (Figure 3D-F), suggesting that transgenic rice expressing *Bs2* together with its NRC helper NLRs suppresses growth of *Xoc* in an AvrBs2-dependent manner.

**Figure 3.**
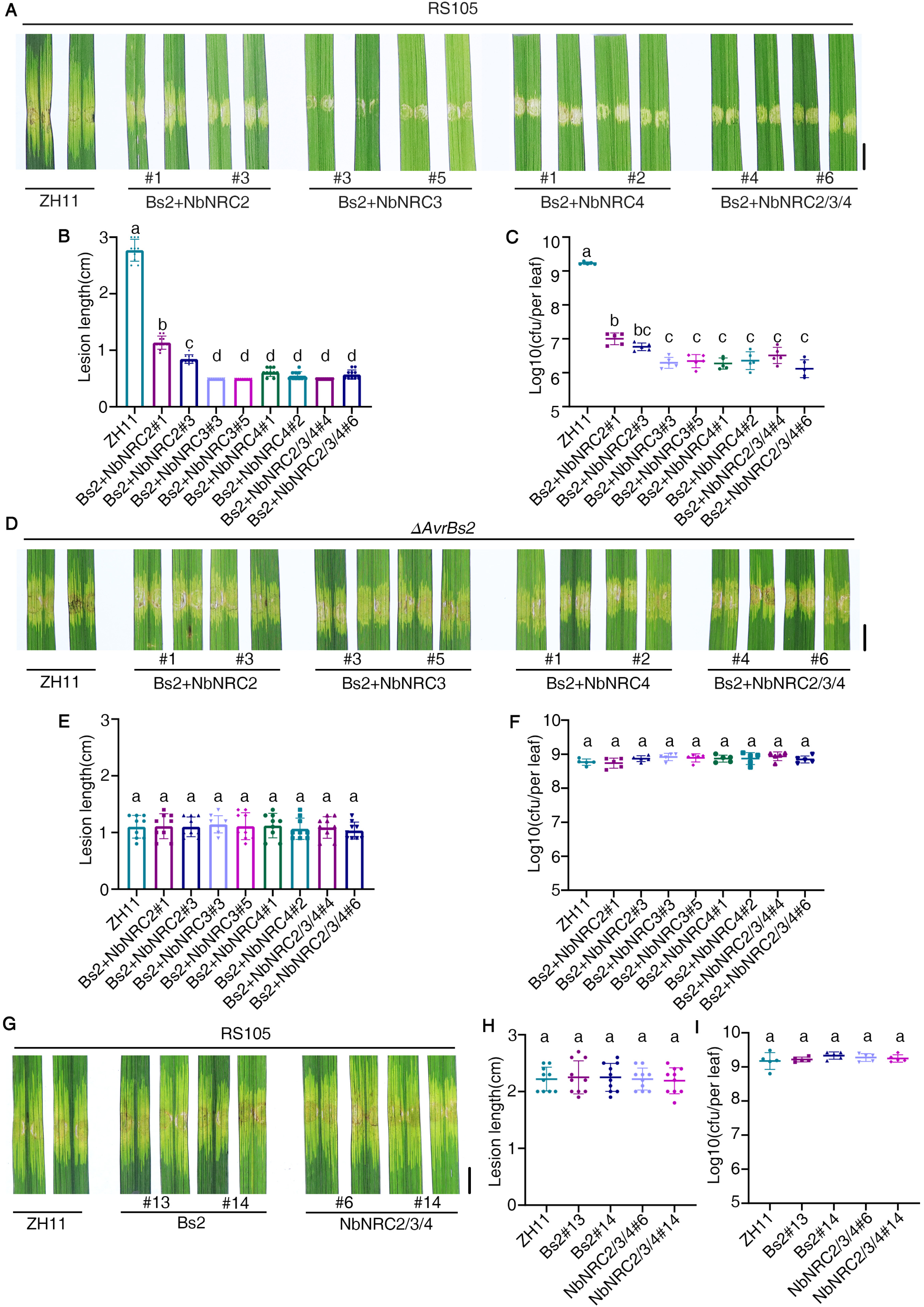
Transgenic co-expression of *Bs2* with NbNRC2, NbNRC3, or NbNRC4 in rice plants confers increased disease resistance against *Xoc*. **(A)** Representative disease symptoms after *Xoc* inoculation. Fully expanded leaves of five-week-old ZH11 and transgenic rice plants co-expressing Bs2 with NbNRC2, NbNRC3 or NbNRC4 were infiltrated with *Xoc* strain RS105. Image of lesion expansion were taken at 14 dpi using syringe-infiltration inoculation. Scale bar, 0.5 cm. **(B)** Quantification of disease lesion length caused by *Xoc* in the wild-type ZH11 and transgenic rice plants co-expressing Bs2 with NbNRC2, NbNRC3 or NbNRC4. Lesion length was measured at 14 dpi. Mean values and standard deviations were calculated from measurements of 10 independent leaves. Statistical significance is indicated by letters (p<0.01, one-way ANOVA followed by Tukey’s post hoc test). This experiment was repeated three times with similar results. **(C)** Bacterial growth of *Xoc* in the wild-type and transgenic lines after inoculation. Leaves of wild-type ZH11 and transgenic lines co-expressing Bs2 with NbNRC2, NbNRC3 or NbNRC4 were infiltrated with *Xoc* strain RS105 carrying AvrBs2. Bacterial titers of a 3-cm leaf segment around the infiltration site were determined at 6 dpi. Statistical significance is indicated by different letters (P < 0.01, one-way ANOVA followed by Tukey’s post hoc test). **(D)** Representative disease symptoms after inoculation with *Xoc* AvrBs2 deletion mutant strain (Δ*AvrBs2*). Fully expanded leaves of five-week-old ZH11 and transgenic rice plants co-expressing Bs2 with NbNRC2, NbNRC3 or NbNRC4 were infiltrated with Δ*AvrBs2*. Image of lesion expansion were taken at 14 dpi using syringe-infiltration inoculation. Scale bar, 0.5 cm. **(E)** Quantification of disease lesion length caused by inoculation with *Xoc* Δ*AvrBs2* mutant strain in the ZH11 and transgenic rice plants co-expressing Bs2 with NbNRC2, NbNRC3 or NbNRC4. Lesion length was measured at 14 dpi. Mean values and standard deviations were calculated from measurements of 10 independent leaves. Statistical significance is indicated by letters (p<0.01, one-way ANOVA followed by Tukey’s post hoc test). This experiment was repeated three times with similar results. **(F)** Bacterial growth of *Xoc* Δ*AvrBs2* mutant strain in the wild-type and transgenic lines after inoculation. Leaves of ZH11 and transgenic lines co-expressing Bs2 with NbNRC2, NbNRC3 or NbNRC4 were infiltrated with *Xoc* Δ*AvrBs2* mutant strain. Bacterial titers of a 3-cm leaf segment around the infiltration site were determined at 6 dpi. Statistical significance is indicated by different letters (P < 0.01, one-way ANOVA followed by Tukey’s post hoc test). **(G)** Representative disease symptoms after *Xoc* inoculation. Fully expanded leaves of five-week-old ZH11 and transgenic rice plants expressing either Bs2 or triple combinations of NRC2/3/4 were infiltrated with *Xoc* strain RS105. Image of lesion expansion were taken at 14 dpi using syringe-infiltration inoculation. Scale bar, 0.5 cm. **(H)** Quantification of disease lesion length caused by *Xoc* in the wild-type ZH11 and transgenic lines expressing either Bs2 or triple combinations of NRC2/3/4. Lesion length was measured at 14 dpi. Mean values and standard deviations were calculated from measurements of 10 independent leaves. Statistical significance is indicated by letters (p < 0.01, one-way ANOVA followed by Tukey’s post hoc test). This experiment was repeated three times with similar results. **(I)** Bacterial growth of *Xoc* in the wild-type and transgenic lines only expressing Bs2 or NRCs after inoculation. Leaves of ZH11 and transgenic lines were infiltrated with *Xoc* strain RS105. Bacterial titers of a 3-cm leaf segment around the infiltration site were determined at 6 dpi. Statistical significance is indicated by different letters (P < 0.01, one-way ANOVA followed by Tukey’s post hoc test).

As a negative control, we generated transgenic rice expressing only *Bs2* or *NbNRCs* (Figure S3E-F). However, transgenic rice expressing either *Bs2* or the combination of the NRC-helpers *NbNRC2/3/4* alone did not exhibit any resistance against *Xoc* (Figure 3G-I), suggesting that transfer of sensor NLRs to recipient species that lack corresponding helper NLRs will be ineffective and helper NLRs are not sufficient to confer resistance. Furthermore, we investigated whether an intact N-terminal MADA motif of NRCs, a conserved molecular signature implicated in resistosome-mediated immune responses(Adachi et al., 2019a; Contreras et al., 2023c), is required for the disease resistance when heterologously expressed. We found that co-expressing of Bs2 with NbNRC2 MADA mutant in which the conserved leucine residues (L9, L13, L17) of NRC2 were substituted by glutamates (NRC2^EEE^) failed to confer resistance (Figure S3 G-J), implying the importance of a functional N-terminal MADA motif of NRC helper in conferring disease resistance against *Xoc* in rice, consistent with the observation that an intact N-terminal MADA motif is required for mediating disease resistance in Solanaceous species(Adachi et al., 2019a; Kourelis et al., 2022).

### Heterologous expression of stacked sensor and helper NLRs in rice does not affect basal resistance, growth and agronomic traits

Since stacking of multiple NLRs in a single cultivar could induce some autoimmunity that effects plant growth and yield (Peng et al., 2021; Zhao et al., 2022), we investigated whether transgenic rice expressing stacked sensor and helper NLRs-encoding genes carry negative effects on plant growth or yield. Phenotypic analysis showed that transgenic rice lines co-expressing *Bs2* with *NbNRC2*, *NbNRC3* or *NbNRC4* individually or in *NbNRC2/3/4* combinations grew normally in the field and exhibited indistinguishable phenotypes to the wild-type ZH11 (Figure 4A). No significant differences were found for major agronomic traits, including plant height, tiller number per plant, grain number per panicle, grain shape, and thousand-grain weight between the transgenic rice lines and ZH11 (Figure 4B-E, and Figure S4A-C). We additionally tested the stack of four NLRs transgenic lines for paddy trials that confirmed co-expressing *Bs2* and *NbNRCs* provides complete field resistance to *Xoc* without affecting growth phenotype in the field (Figure S4D-F). We also detected no significant difference in genome-wide transcriptional landscapes between these transgenic plants and ZH11 plants under unchallenged conditions (Figure S4G), again demonstrating no sign of constitutive immunity. Thus, stacking sensor-helper NLRs could confer disease resistance without resulting in autoimmunity or negative effects on agronomic traits, in contrast to the negative effects occasionally observed for multiple sensor NLRs stacking (Peng et al., 2021).

**Figure 4.**
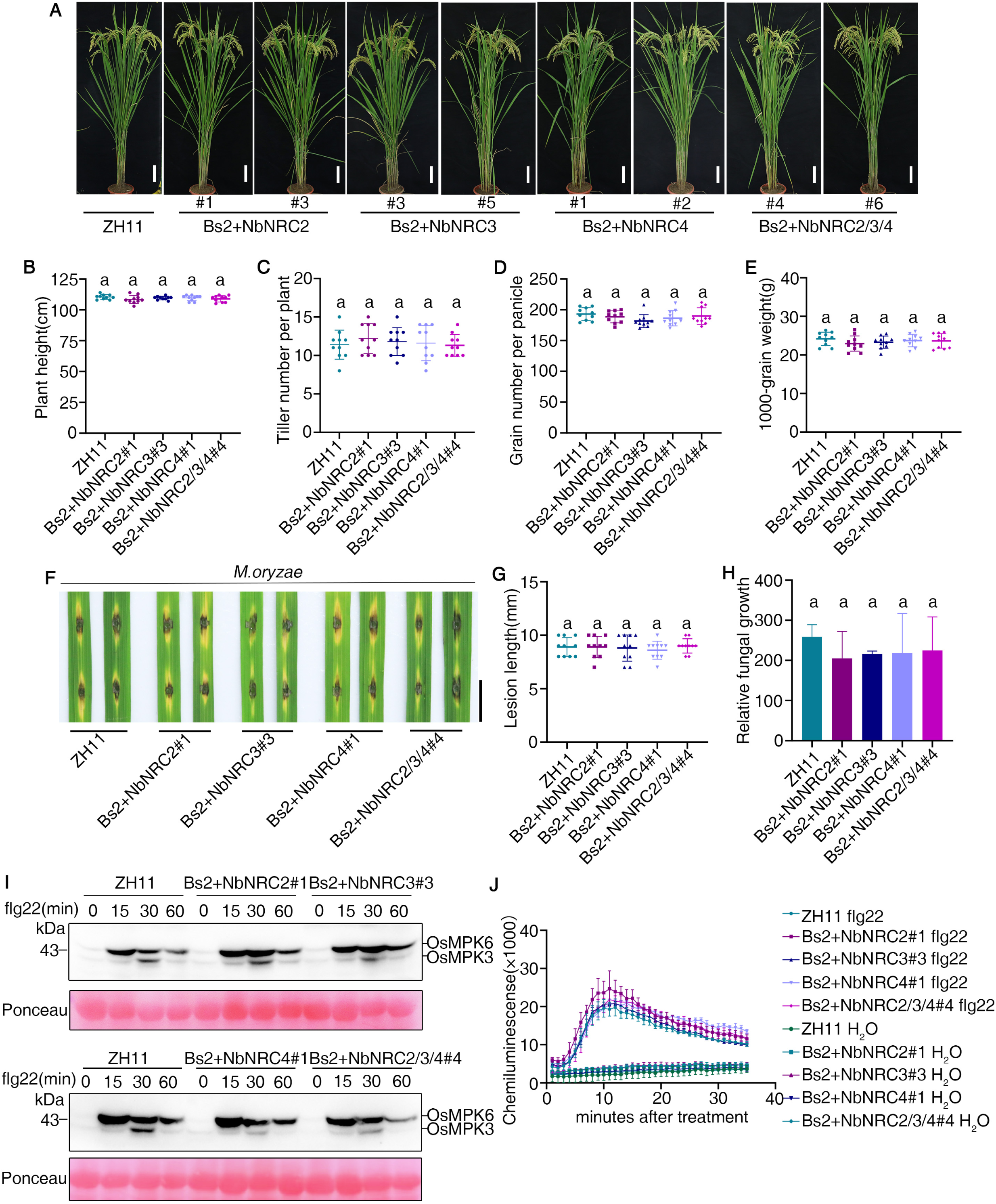
Comparisons of growth phenotypes, agronomic traits and basal immunity between wild-type ZH11 and transgenic lines coexpressing Bs2 with NbNRCs. **(A)** Comparisons of growth phenotypes between wild-type ZH11 and transgenic lines co-expressing Bs2 with NbNRCs. ZH11 and transgenic plants were grown in the field and photographed at tillering stage. Scale bar, 10cm. **(B)-(E)** Evaluation of agronomic traits of transgenic rice lines co-expressing Bs2 with NbNRCs and ZH11 plants in the field. The average plant height (B), total tiller number per plant (C), grain number per panicle (D), and thousand-grain weight (E) of ZH11 and transgenic lines are shown, respectively. **(F)-(H)** The disease symptoms and relative fungal biomass of representative leaves of transgenic lines co-expressing Bs2 with NbNRCs and ZH11 control. Wild-type rice ZH11 and transgenic lines co-expressing Bs2 with NbNRCs were inoculated with *M. oryzae* isolate RB22 by punch inoculation. Punch-inoculated plants were recorded 7 dpi, respectively. Quantification of lesion lengths and fungal biomass of punch inoculation is shown in (G) and (H), respectively. Values are means ± SD, n = 3 (technical repeats). Data were analyzed by one-way ANOVA followed by Tukey test. Both inoculations have been replicated three times with similar results. **(I)** Flg22-induced MAPKs activation profile in the wild-type control ZH11 and transgenic seedlings co-expressing Bs2 with NbNRCs. Ten-day-old seedlings were treated with 10 µM flg22 and collected at the indicated time points. MAPK activation was analyzed by immunoblot analysis using phosphor-p44/42 MAPK (Erk1/2) (top panel), and the equal protein loading is indicated by Ponceau S staining for Rubisco (RBC) (bottom panel). **(J)** ROS generation induced by flg22 treatment in transgenic lines co-expressing Bs2 with NbNRCs and wild-type control ZH11. Leaf discs were treated with 10 µM flg22. ROS generation was monitored for 30 min. Values are means ± SD calculated from at least three technical repeats.

To examine whether BLS resistance is related to the elevated basal resistance that may arise from ectopic expression of multiple NLRs, transgenic rice lines co-expressing *Bs2* with *NbNRC2*, *NbNRC3* or *NbNRC4* individually or in combinations were challenged with a virulent strain of *M. oryzae* for basal immunity evaluation.

These transgenic plants showed no significant difference in resistance with similar lesions and fungal biomass compared with the ZH11 (Figure 4F-H), indicating similar level of rice blast susceptibility. In addition, we also checked whether basal resistance markers including MAPK activation and reactive oxygen species (ROS) burst upon PAMP perception are altered in the transgenic lines. Both ZH11 and transgenic lines exhibited comparable levels of ROS production and activation of MAPKs upon flg22 or chitin treatment (Figure 4I-J and Figure S4H-I). In conclusion, the basal resistance of transgenic lines is comparable with wild-type ZH11 control plants. Therefore, cross-species transfer of sensor and helper NLR stacks does not produce symptoms of autoimmunity and compromise plant fitness.

To further investigate the enhanced resistance phenotype of transgenic rice, we measured the expression levels of defense-related marker genes during *Xoc* infection. Expression levels of defense-related marker genes such as *OsPR1a*, *OsPBZ1*, and *OsKSL4* in transgenic and ZH11 plants were similar before inoculation. However, these genes were significantly upregulated in transgenic rice after *Xoc* inoculation while such induction of defense-related genes was not observed in the ZH11 controls (Figure S4J), suggesting that *Bs2* and *NbNRCs* co-expression resulted in strong induction of defense-related gene expression in response to *Xoc*. We further analyzed genome-wide changes in transcriptional processes between transgenic and ZH11 plants. Transgenic rice co-expressing *Bs2* and *NbNRC2/3/4* were chosen for transcriptomic comparisons since we had previously performed RNA-seq under unchallenged conditions. The infiltrated leaf parts were collected at 0h and 24h after inoculation with *Xoc* for transcriptome profiling. We identified 994 differentially expressed genes (DEGs) (p<0.05, fold change>2) in the transgenic lines compared with ZH11 controls (Figure S4K and Supplemental Table 1). When compared with public available transcriptome changes induced by NLRs activation such as Pi9, a rice blast NLR that activates immune responses upon recognizing *M. oryzae* avirulence effector AvrPi9, and PRRs activation such as OsFLS2, a PRR triggering immune activation upon sensing of flg22(Jain et al., 2017; Tang et al., 2021). Bs2- and NRC-mediated immune responses showed a large overlap (266/994, 26.7%) with Pi9-induced responses but considerably less overlap (40/994, 4%) with flg22-induced responses (Figure S4L). Moreover, Bs2- and NRC-induced DEGs contain 253 Pib-specific DEGs but only 26 flg22-specific DEGs. These data suggest that like Pi9, NLR-activated transcriptome reprogramming is activated in transgenic lines upon *Xoc* infection, reinforcing the concept that sensor and helper NLRs are functionally transferable and reconstitute an immune signal transduction module in the recipient species.

## Discussion

The discovery of NRC helper NLRs has transformed our understanding of immune receptor mechanisms in the asterid clade of angiosperms (Wu et al., 2017). Upon effector recognition, activated sensor NLRs via their NB domains activate NRC helpers to form hexameric resistosomes that do not incorporate the sensor NLR (Ahn et al., 2023; Contreras et al., 2024; Contreras et al., 2023c; Selvaraj et al., 2024). NRC clade helper NLRs are necessary for cognate sensor NLRs to activate defense. We investigated whether NRC helpers are sufficient to support sensor NLR function when co-delivered to non-asterid plants, or whether additional components might be required. We showed that in soybean, Arabidopsis and rice, co-delivery of sensor NLRs with cognate NRC helper NLRs is sufficient to enable sensor NLR function and to break RTF. Compared to NLRs, PRRs are more evolutionarily ancient immune receptors and more likely to share signaling pathways between all angiosperms (Yue et al., 2012). Indeed, the Arabidopsis EFR receptor can function in rice (Lu et al., 2015; Schwessinger et al., 2015). Thus, this approach has particular relevance for intracellular rather than cell surface immune receptors.

The interfamily utilization of sensor and helper NLR genes would overcome RTF, thus broadening the resistance utility of known NLRs and enabling breeders to redeploy known NLR genes across different plant families. These insights have many potential uses but we highlight one. The bacterial pathogen *Xoc* causes major disease losses in rice. Breeding for disease resistance is preferable to chemical control, but no genes for *Xoc* resistance are available in the rice gene pool. We show here that pepper NLR immune receptor Bs2 can confer BLS resistance when co-expressed in rice with the appropriate NRC helper NLRs. The pepper NRC-dependent Bs2 receptor confers responsiveness to AvrBs2*^Xoc^* as well as AvrBs2*^Xcv^*. Transgenic rice co-expressing *Bs2* with *NbNRC3* or *NbNRC4* acquire the ability to respond to AvrBs2*^Xoc^* and become fully resistant to *Xoc* infection. Previous attempts to transfer *Bs2* to phylogenetically distant Arabidopsis and cassava were limited by RTF (Díaz-Tatis et al., 2019; Tai et al., 1999), likely due to the requirement for NRC helper NLRs. Our findings suggest that asterid NLR repertoires can now be deployed to elevate disease resistance in any angiosperm. Interestingly, some NRCs partner better with Bs2 than others; co-expressing *Bs2* with *NbNRC2* results in moderate resistance compared to using *NbNRC3* or *NbNRC4*. We do not exclude the possibility of another unknown *Xoc* effector which can restrict the activity of NbNRC2 but not NbNRC3 and NbNRC4 (Contreras et al., 2023b; Derevnina et al., 2021).

Bs2+NRCs rice displays NLR-induced transcriptome features upon *Xoc* infection. Comparative transcriptomic analysis showed *Bs2* and *NRCs* mediate genome-wide transcriptional reprogramming in response to *Xoc* inoculation similar to the ETI-associated transcriptome caused by NLR genes conferring blast resistance upon *M. oryzae* infection. The induction of the rice ETI-associated transcriptome is correlated with disease resistance against *Xoc*.

Heterologous overexpression of multiple NLRs in plants can lead to constitutive activation of immune responses, resulting in necrosis and/or yield penalties (Peng et al., 2021; Zhao et al., 2022). In our study, transgenic rice co-expressing *Bs2* and *NRCs* did not show such a phenotype. No constitutive production of ROS, activation of defense-related gene expression or growth defects or adverse fitness consequences were observed in the field. Moreover, transgenic rice co-expressing *Bs2* and *NRCs* were not more resistant to the fungal pathogen *M. oryzae*. Therefore, Bs2+NRC co-expression does not constitutively activate defense responses.

In conclusion, interfamily co-transfer of Bs2 and NRC helper NLRs, but not either alone, endows rice with robust resistance against *Xoc*, demonstrating the feasibility of breaking RTF. Our finding also provide a paradigm for controlling diseases caused by evolutionarily related pathogens through redeploying a known NLR that is able to recognize orthologous effectors from closely related pathogen species. Some *P. infestans* effectors are conserved across multiple *Phytophthora* species, and previously reported Solanum NLRs that recognize *P. infestans* effectors are also able to recognize orthologous effectors from multiple *Phytophthora* species (Oh et al., 2024). Conceivably, co-transferring these Solanum NLRs together with NRCs could provide non-Asterid crops such as soybean, Brassicas, cacao and strawberry with resistance against other *Phytophthora* species. This suggests that NLRs derived from more distantly related species have enormous potential as untapped genetic resources to expand the disease resistance gene pool available for improving disease resistance in crops. We speculate that this strategy could potentially provide more durable resistance since NLRs derived from distantly related plant species could be less susceptible to effector-mediated suppression by non-adapted pathogens than NLRs from the host plants (Oh et al., 2023).

## Acknowledgments

This research was financially supported by the National Natural Science Foundation of China (Grant No. 32270307), the National Key Research and Development Project (2022YFD1401400), Beijing NOVA Programme (Z211100002121066), the Gatsby Charitable Foundation (UK) and the Pinduoduo-China Agricultural University Research Fund (Grant No. PC2023B02014). We thank Prof. Sophien Kamoun for sharing plasmids and *nrc2/3/4 N. benthamiana* CRISPR seeds and helpful discussions. We thank Dr. Hee-Kyung Ahn and Dr. Sarah Pottinger providing assistance with Arabidopsis transgenic lines generation and validation.

## Materials and methods

### Plant materials, bacterial strains and growth conditions

Arabidopsis plants were grown in a controlled environment room (CER) at 22°C under an 8-h light/16-h dark cycle. For *Pseudomonas* infiltrations, 4-5 week old plants were used. *N. benthamiana nrc2/3/4* CRISPR lines have been described previously (Contreras et al., 2023c). *N. benthamiana* wild-type and *nrc2/3/4* CRISPR plants were grown in a CER at 22°C and 45%–65% humidity under a 16-h light/8-h dark cycle. For *Agrobacterium* infiltrations, 4-5 week old plants were used. The rice variety Zhonghua 11 (*Oryza sativa* subsp. *japonica*) and transgenic plants were grown in the greenhouse with 16 h day and 8 h night cycle at 24-28°C. *X. oryzae* pv. *oryzicola* strain RS105 were grown in NA medium containing rifampin (50 μg/mL) at 28°C. For agronomic traits evaluation, all of the tested lines were planted at the Shangzhuang experimental farm of China Agricultural University in Beijing. For pathogen inoculation, nontransgenic controls and transgenic rice plants were grown in the greenhouse with 16 h day and 8 h night cycle at 24-28°C.

### Plasmid construction, generation and identification of transgenic rice plants

Constructs in this study were generated using Golden Gate cloning (Engler et al., 2008), which enables assembly of multiple DNA fragments into a destination vector. The CDS of sensor and helper NLRs were assembled into golden gate level 0 entry vector pICSL01005. For constructs used for rice transformation, level 0 modules containing Bs2, NbNRC2, NbNRC3 and NbNRC4 were used as described previously (Contreras et al., 2023c; Wu et al., 2017). The level 0 constructs were assembled with different promoters, terminators and C-terminal epitope tags into level 1 vectors at different positions. The level 1 units were assembled in level 2 acceptor pICSL4723 along with a hygromycin resistance as a plant selectable marker.

For constructs used for Arabidopsis transformation, the sensor NLRs were then assembled in a level 1 vector (pICH47761) with the Arabidopsis Ubiquitin10 promoter (pICSL12015), a C-terminal epitope tag 3XFLAG (pICSL50007) and a Nos terminator (pICH41421). Rpi-amr3 cloning in a level 0 vector has been described previously (Lin et al., 2022); Bs2 (pICSL80119) was used in level 1 construct assembly. The helper NLRs were assembled in a level 1 vector (pICH47772) with a 35S promoter (pICH51277), a C-terminal epitope tag 4XMYC (pICSL50010) and an Ocs terminator (pICH41432). The sensor-helper or sensor alone constructs were then assembled in a level M vector (pAGM8031) along with an NPTII selection marker (pICSL11055), FastRed (pICSL11039) and a small linker (position 3-pICH50433). The level M stack was subsequently assembled in a level P vector, pRKan290 (GenBank M61152). The level P plasmid was then transformed into Agrobacterium strain AGL1 for transient assays and plant transformations. AvrAmr3 and AvrBs2 used for agroinfiltration assays have been described previously (Contreras et al., 2023c; Lin et al., 2022).

For constructs used for HR assay in Arabidopsis, AvrAmr3^59-339^ and AvrBs2^62-714^ were assembled into golden gate level 0 entry vector pICSL01005. AvrAmr3^59-339^ derivative that lacks the signal peptide and the RXLR motif has been described previously (Lin et al., 2022). AvrBs2^62-714^ derivative that lacks the secretion signal has been previously described to function in planta (Wei et al., 2000). The effectors were then assembled into a golden gate compatible version of pEDV3 (Sohn et al., 2007) along with a C-terminal epitope tag V5 (pICSL50012). pEDV3 contains the 136 amino acids that encode a T3SS and the AvrRPS4 promoter. The assembled effector construct was electroporated (Choi et al., 2006) in the effector-less mutant of *Pseudomonas syringae* pv. *tomato* DC3000, D36E (*Pst* DC3000 D36E).

For rice transformation, the *japonica* rice cultivars, Zhonghua 11 (ZH11) were used as the recipients. The positively transformed plants were identified by selecting hygromycin-resistant seedlings and checking the expression of transgenes by immunoblot analysis. For Arabidopsis transformation, *Arabidopsis thaliana* Col-0 was used for making stable transgenic. For generation of stable lines, floral dip transformation was used (Clough and Bent, 1998). Transgenic T1 progeny was selected through screening red seeds (FastRed selection). The T1 seeds were then grown for generating T2 seeds which were used in the cell death assay.

### RNA extraction and quantitative real-time PCR

Total RNA was extracted from collected leaf segments using TRIzol Reagent (Invitrogen). To synthesize cDNA, 1 μg of RNA was used in a 20 μL reaction volume using a First-strand cDNA Transcriptase kit (TransGen) following the manufacturer’s instruction. qRT-PCR was performed on ABI QuantStudio 6 Flex Real-Time PCR System using the SYBR Premix ExTaq kit (Takara) in accordance with the manufacturer’s instructions. The gene-specific primers used for qRT-PCR analysis are listed in Supplementary Table 2. The rice Ubiquitin (*OsUbq*) gene was used as an internal control.

### RNA-seq analysis

For RNA-seq experiments, about 0.2g leaves infected with *Xoc* or MgCl_2_ were collected at 24 hpi, and three biological replicates were collected for each treatment. Total RNA was isolated using the Trizol Reagent (Invitrogen Life Technologies), after which the concentration, quality and integrity were determined using a NanoDrop spectrophotometer (Thermo Scientific). Three micrograms of RNA were used as input material for library preparation and sequencing. The sequencing library was then sequenced on NovaSeq 6000 platform (Illumina) Shanghai Personal Biotechnology Cp. Ltd. The reference genome and gene annotation files were downloaded from genome website (IRGSP-1.0). Then difference expression of genes was analyzed by DESeq (v1.38.3) with screened conditions as follows: expression difference multiple log2(FoldChange)> 1, significant P-value<0.05.

### Protein extraction and immunoblot analysis

Protein samples were prepared from rice leaf segments and were homogenised in extraction buffer (125 mM Tris-HCl, pH 8.8, 1% (w/v) SDS, 10% (v/v) glycerol, 50 mM Na_2_S_2_O_5_). The supernatant obtained after centrifugation at 12,000g for 10 min was separated by SDS-PAGE. After transferring proteins from gels to polyvinylidene fluoride (PVDF) membranes (Merck-Millipore), membranes were probed with horseradish peroxidase (HRP)-conjugated FLAG antibody (A5892, Sigma), HRP-conjugated HA antibody (12013819001, Merck-Roche), or V5 antibody (AE017, ABclonal). Anti-mouse IgG (whole molecule)–peroxidase antibody produced in goat (AS003, ABclonal) was used as secondary antibody following the use of V5 antibody. The protein bands were detected by luminescence of HRP using the chemiluminescent substrate on a Tanon-5200 Chemiluminescent Imaging System (Tanon Science and Technology).

For confirming the expression of sensor-helper NLR and sensor NLR alone constructs expressed transiently in *N. benthamiana* or in stable Arabidopsis transgenic lines, total protein was extracted from the plant tissue as described previously with slight modifications (Ahn et al., 2023). Briefly, 0.1 gram (g) of leaf tissue homogenized in 200ul of extraction buffer (50□mM Tris-HCl pH 7.5, 50□mM NaCl, 50mM MgCl_2_, 10% Glycerol, 10□mM DTT, 0.2% NP-40, protease inhibitor cocktail). The samples were then centrifuged at 12,000g for 20 min at 4°C to remove the debris. The resulting collected supernatant was mixed with 3XSDS PAGE loading dye. The samples were then heated at 65°C for 15 min. The extracted samples were resolved on 10% (w/v) SDS-PAGE gels and were subsequently transferred onto PVDF membranes with the TransBlot (Biorad). The membrane was probed with 1:10000 dilution of either anti-FLAG (Sigma, A8592) for detecting sensor NLRs or with anti-MYC (Sigma, 16-213) antibody for detecting helper NLRs. For protein detecting, the blot was probed with ECL substrates (Thermo Fisher, 34580).

### LUC activity assay

For rice protoplasts transient assay, rice protoplasts were isolated from rice etiolated seedlings as described previously. The firefly luciferase gene (LUC) driven by the ubiquitin promoter serves as a reporter to monitor protoplasts viability. The combinations of the indicated constructs were co-transfected together with the LUC plasmid into rice protoplasts using PEG-calcium-mediated transformation. Luciferase activity was measured 24h after transfection using a luciferase assay system (Promega) following the manufacturer’s instructions, and the reduction in luminescence was compared with protoplasts expressing the control GFP vector.

For soybean protoplasts transient assay, soybean protoplasts were isolated using the method outlined in Xiong et al. (2019) with a few modifications. Soybean cultivar Williams 82 was grown in a growth chamber under a 16-hour photoperiod at 28°C for two weeks, then etiolated in the dark for four additional days. The first fully expanded trifoliate leaf was harvested and cut into thin strips 1-2□mm thick. Two grams of leaf material were added to 40□mL of digestion solution and left to digest for five hours on a gyratory shaker at 60□rpm. The protoplasts were quantified and diluted to a concentration of 2□× 10^6^□cells/mL and transfected using a method adapted from Yoo et al. (2013) and Xiong et al. (2019), scaled up for high-throughput format. A total of 1.5□×□10^5^ protoplasts were transfected per well of a 2.2□mL 96-deep-well plate with plasmid DNA at a concentration of 2□µg/µL, totaling 28□µg per well. As outlined in Saur et al. (2019), the plasmid DNA consisted of Luciferase, NLR, Avr, and Helper genes at a ratio of 5:3:3:3, all under the control of a double 35S promoter, with a 3×FLAG C-terminal tag and a rbcS terminator. The plate of transfected protoplasts was left in the dark for 2-3 days before analysis, following the methodology of Saur et al. (2019), using Promega cell lysis solution (catalogue number E1531) and luciferin (catalogue number E1501), in a standard white 96-well plate. Measurements were taken using a ThermoFisher Scientific Varioskan LUX microplate reader. The plate was measured for one second per well, with three readings per well.

### Magnaporthe oryzae and Xoc inoculation assays

For *Xoc* inoculation, leaves of 3-week-old plants were infiltrated with a bacterial suspension using a 1 mL syringe without needle at an optical density of 600 nm (OD_600_) of 0.3, and symptoms of water-soaked lesions were scored 14 days post-inoculation (14 dpi). For *Xoc in planta* growth assay, leaves were infiltrated with a bacterial suspension at a final OD_600_ of 0.3, and a segment of 3 cm around the infiltration site was collected at 7 dpi.

For *M. oryzae* punch inoculation, *M. oryzae* strain RB22 spores were collected in sterile water and the spore concentration in the suspension was adjusted to approximately 5□×□10^5^ conidia per ml. Detached leaves of 6 to 8-week-old rice plants were lightly wounded at two spots on each leaf using a mouse ear clip, the punched sites were then treated with one drop of the spore suspension. The inoculated leaf were kept in a culture dish that contains 0.1% 6-benzylaminopurine (6-BA) and lesion length were measured using at 6 dpi.

### *Pseudomonas* infiltrations

*Pst* DC3000 D36E carrying specific effectors or empty vector was grown on King’s B agar plate supplemented with antibiotics (rifampicin: 50 µg/mL and gentamycin 20 µg/mL) at 28°C for 2 days. Bacteria was harvested from the plates and was resuspended in 10 mM MgCl_2_ to the required OD_600_ of 0.2 (1 x 10^8^ cfu/mL). Leaves of 4-5 week old Arabidopsis were hand infiltrated with a needless syringe and were monitored for symptoms from 2 dpi.

### Agromonas assay

For testing the delivery of effectors by *Pst* DC3000 D36E T3SS, agromonas assay was conducted in *N. benthamiana* as described previously (Buscaill et al., 2021). Briefly, *Agrobacterium* carrying the respective sensor NLR or empty vector was infiltrated in *N. benthamiana*. The infiltrated area was marked on each leaf. Two days after agro-infiltration the *Pst* DC3000 D36E carrying the respective effector or empty vector was infiltrated in marked area. Cell death was observed 3 days post *Pseudomonas* infiltration.

### Agrobacterium-mediated transient HR assays in Nicotiana benthamiana

*Agrobacterium* strain GV3101 carrying various constructs were grown in liquid LB-medium supplemented with the appropriate antibiotic for 24 hr at 28°C and 200rpm at incubator shaker. Cells were harvested by centrifugation, washed in 5 ml of 10 mM MgCl_2_, and re-suspended at OD_600_=0.5 in infiltration medium (10 mM MgCl_2_, 10 mM MES, pH 5.6, 1 mM acetosyringone) for infiltration. Leaves of 4–5 weeks old *N. benthamiana* were infiltrated for HR assay. Agroinfiltrated leaves were harvested for photographing at 4-5 dpi.

### Electrolyte leakage assays

For ion leakage measurements in *N.benthamiana*, five leaf dics (diameter=1cm) per measurement were collected form the infiltrated area at 20 hpi, washed in ddH_2_O for 30 min to remove ion caused by wounding. Leaf discs were carefully transferred to 2.5ml ddH_2_O in 24-well plate. Electrolyte leakage was measured at indicated time points with a conductometer FiveEasy F30 (Mettler Toledo).

For ion leakage measurements in Arabidopsis, ion leakage from leaf discs corresponding to the infiltrated area of Arabidopsis leaves was measured with the aid of a conductivity meter (WTW, Weilheim, Germany) (Alam et al., 2021). Briefly, leaf discs taken from the infiltrated area were submerged in 15ml of water and cell death was quantified at the indicated time points. At least three technical replicates were used for each independent T2 line. Cell death in intact leaves was detected by taking images under a UV light.

## Author contributions

Project conceptualization: H.G., and J.D.G.J.;

Experimental design: H.G., J.D.G.J. and W.S.;

Generation of transgenic rice, inoculation assay and rice part-related experiments: X.D., Y. Z., F.C.;

Generation of transgenic Arabidopsis, inoculation assay and Arabidopsis part-related experiments: M. A.;

Soybean protoplasts transient transformation: K. W., L. M., J. H.;

Bioinformatics analysis: Y. Z.;

Rice field trials: X.D., F.C., W.S;

Writing– original draft: H.G., J.D.G.J. and M. A.;

Writing– review & editing, H.G., J.D.G.J., M. A. and W.S.;

Funding acquisition: H.G., J.D.G.J. and W.S.

## Supplementary Figure legends

**Supplemental Figure S1.**
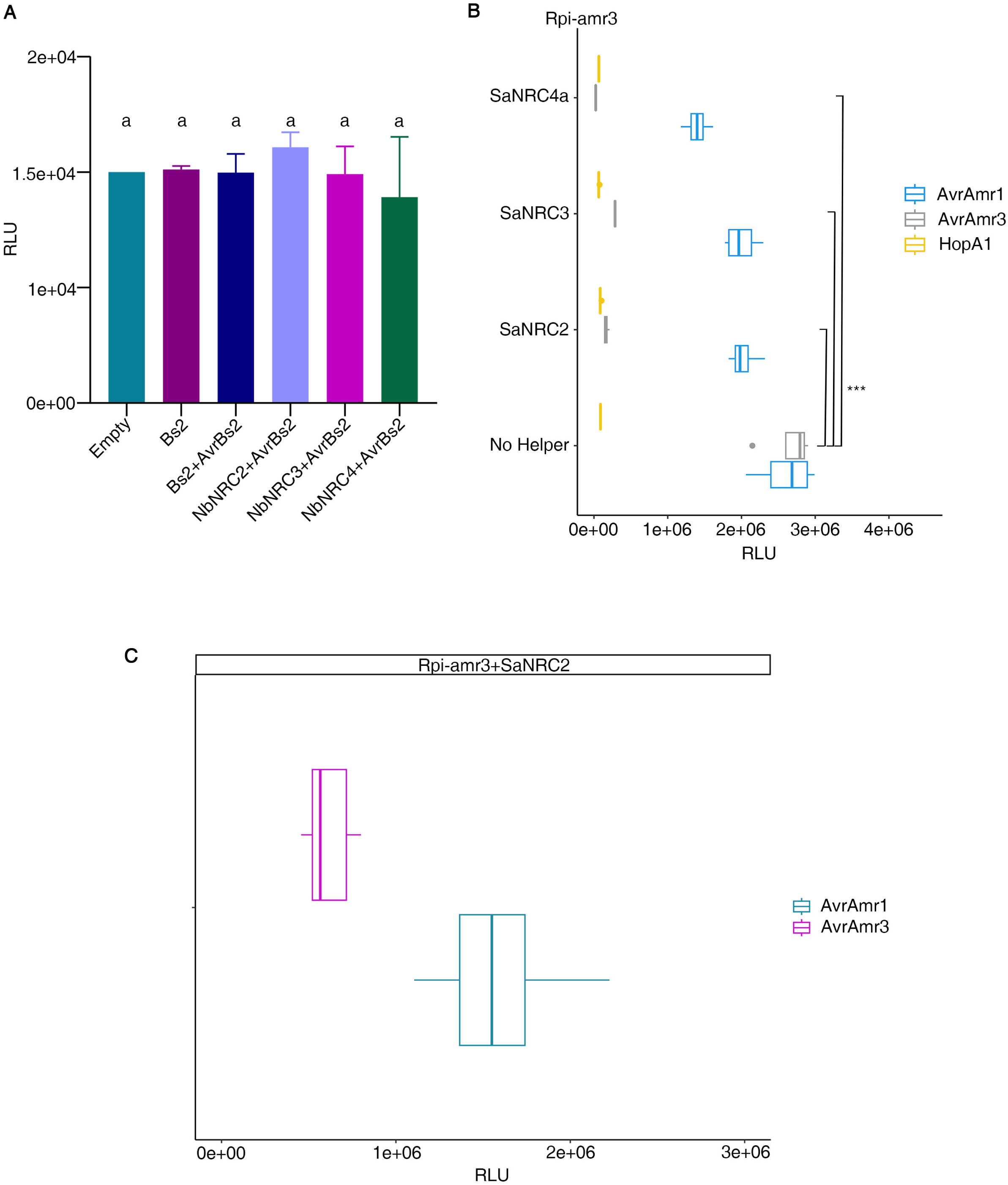
Transient assays verifying sufficiency of NRC helpers to support sensor NLR function in non-asterids. (A) No reduction in luciferase activity upon co-expression of AvrBs2 and NRCs. The indicated constructs were co-transfected with a luciferase reporter vector into isolated rice protoplasts, and luciferase activity was measured 24 h post transfection. The experiment was independently repeated three times with similar results. (B) Co-expression of Rpi-amr3 with AvrAmr1, AvrAmr3 or cell death control HopA1, in the presence or absence of NRCs, in soybean protoplasts. The protoplasts were also transfected with Luciferase reporter. Luciferase activity was measured in relative light units 48 h post transfection. The experiment was repeated twice with similar results. (C) Effector recognition by Rpi-amr3+SaNRC2 results in cell death in Arabidopsis protoplast. Rpi-amr3+SaNRC2 were co-expressed with AvrAmr3 or AvrAmr1 control in Arabidopsis protoplasts that had been transfected with Luciferase reporter. Luciferase activity was measured in relative light units 48 h post transfection. The experiment was repeated independently three times with similar results.

**Supplemental Figure S2:**
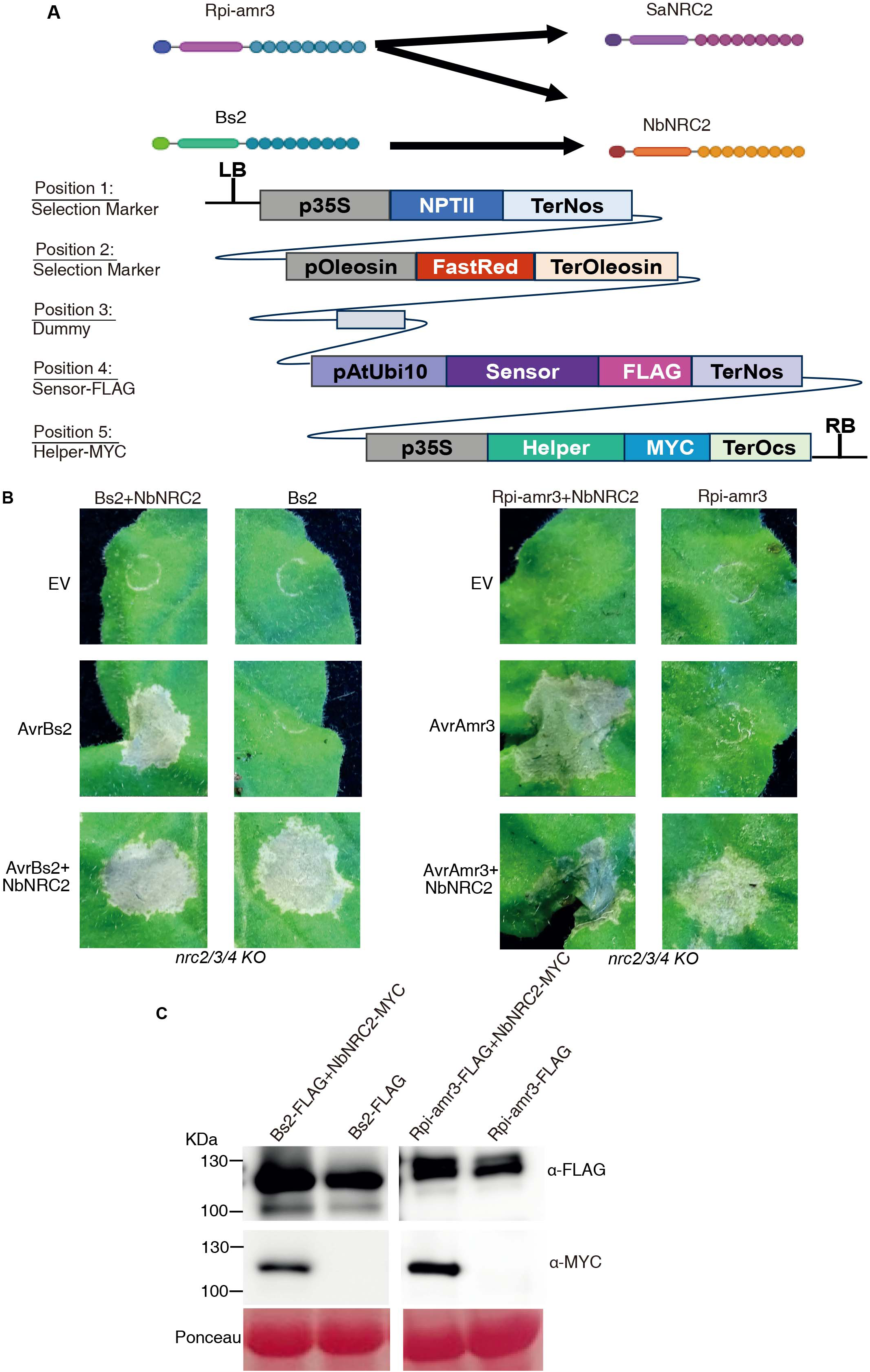
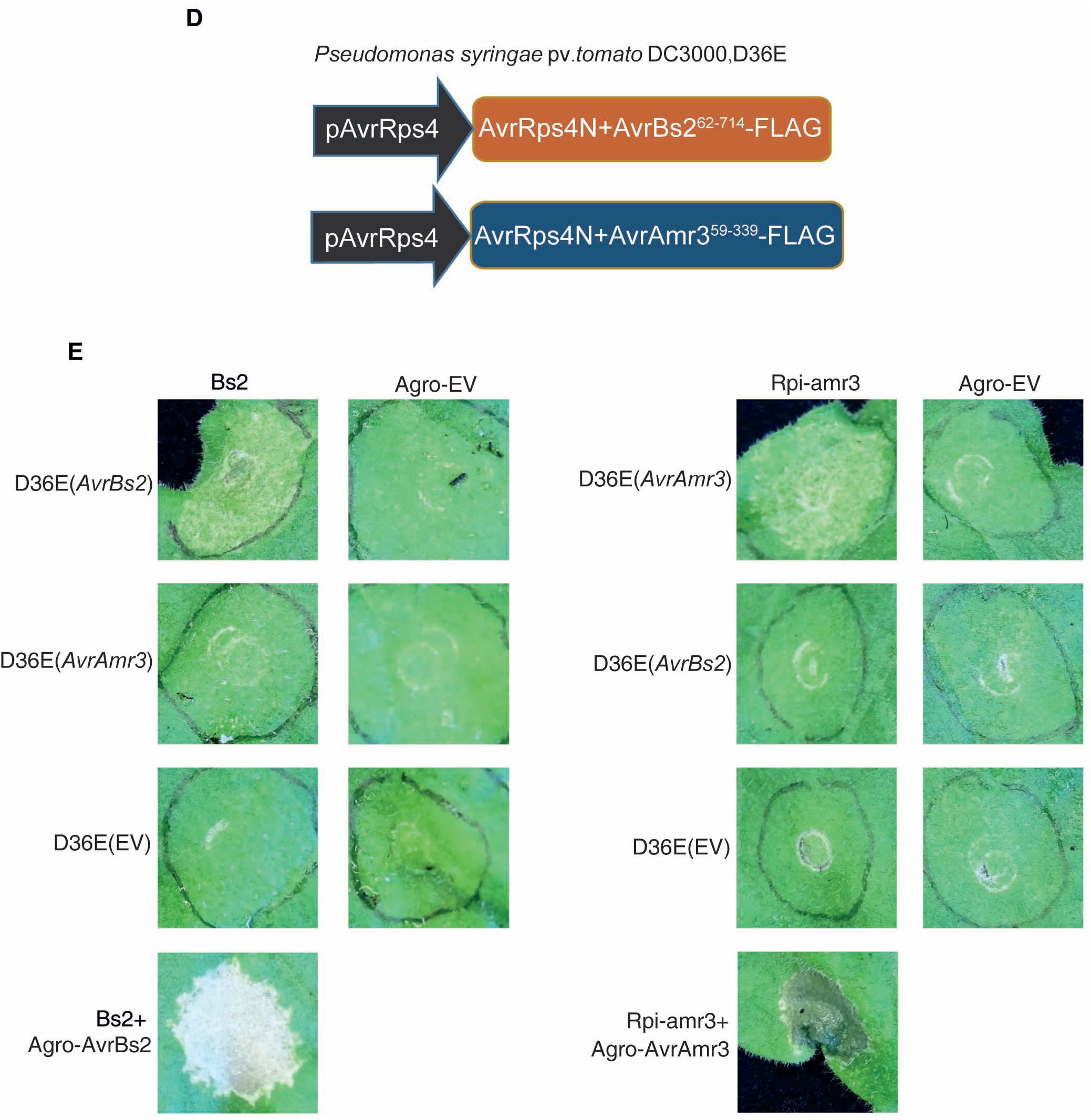
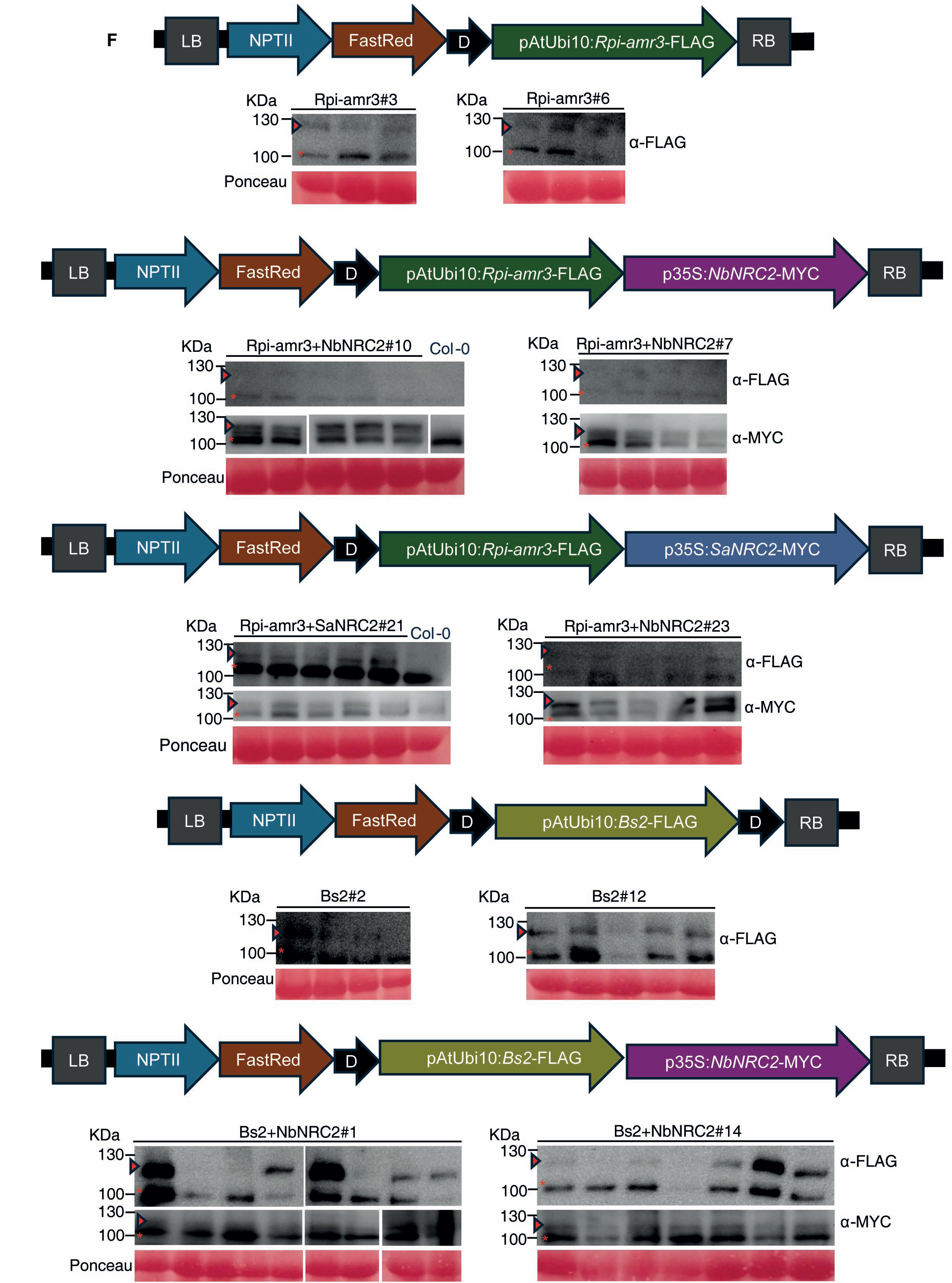
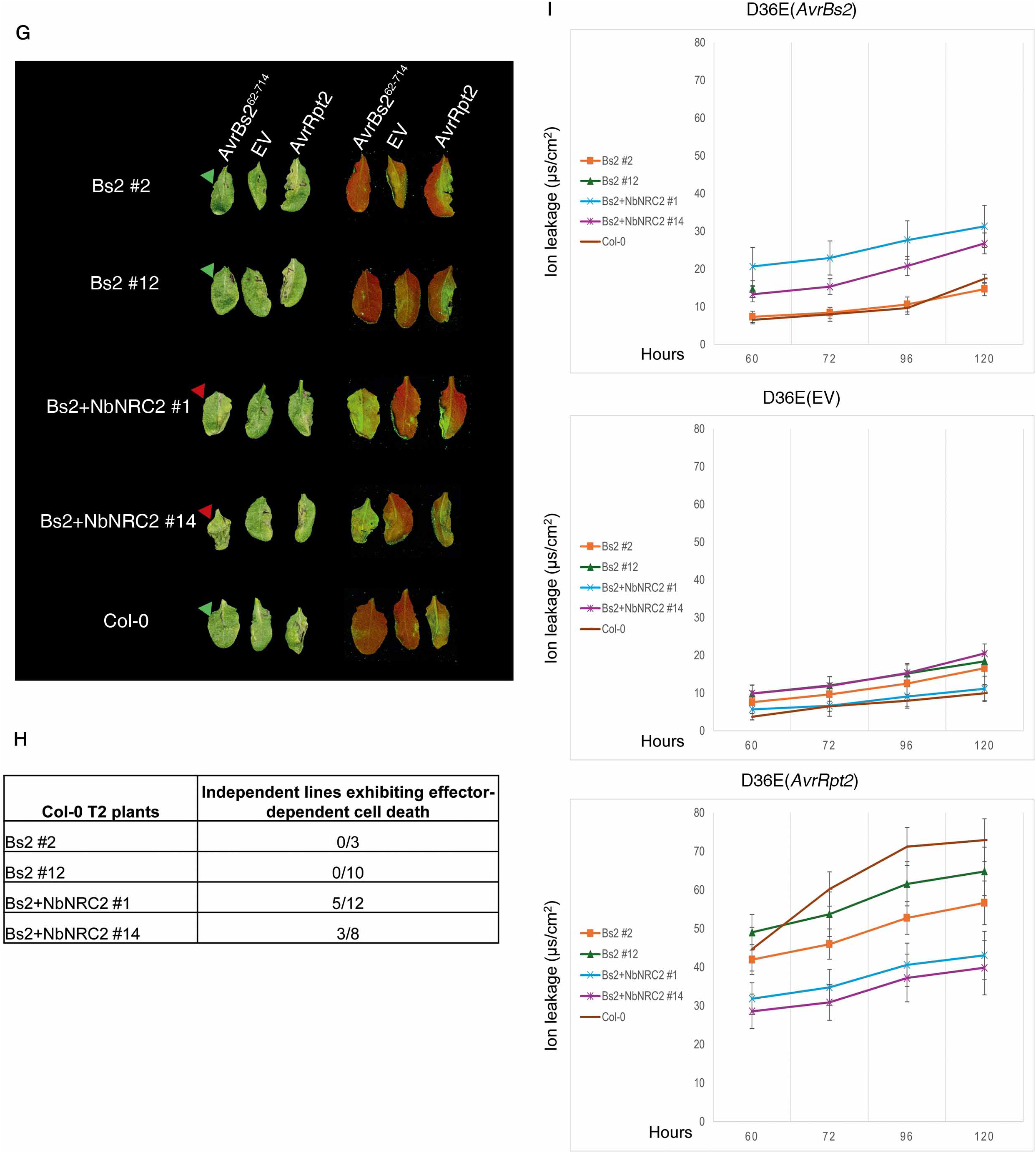
Solanaceae sensor and helper NLRs transformed in Arabidopsis recognize the cognate effector delivered by bacterial T3SS. (A) Construct design for Arabidopsis transformation. Solanaceae sensor NLRs used for Arabidopsis transformation included Rpi-amr3 from *Solanum americanum* and Bs2 from pepper. Solanaceae helper NLRs used for transformation included NRC2 from either *S. americium* or *N. benthamiana*. Each construct was assembled with a promoter and a 3’UTR module. NPTII (position 1) and Fast red (position 2) were used as selection markers. Dummy (position 3) is used as a small linker between the selection markers and the sensor NLRs (position 4). Sensor and helper (position 5) NLRs are epitope tagged at the C-terminus. (B-C) Validation of sensor-helper or sensor alone constructs expression and function when transiently expressed in *N. benthamiana nrc2/3/4* KO plants. (B) Sensor-helper and sensor only constructs were infiltrated (OD_600_=0.5) with either empty vector (EV), respective effector or effector and sensor. Images were taken 4 days post-infiltration (dpi). (C) Anti-FLAG and Anti-MYC immunoblot for sensors and helper respectively shows protein accumulation of different constructs 2 dpi in *N. benthamiana* leaves. Panel below shows Ponceau stain of Rubisco as a loading control. (D-E) Effector delivery using T3SS in *N. benthamiana*. (D) AvrAmr3^59-339^ and AvrBs2^62-714^ were assembled into a golden gate compatible version of pEDV3. pEDV3 contains a T3SS and the AvrRps4 promoter. The effectors were epitope tagged at the C-terminal. (E) Sensor NLRs (Bs2 and Rpi-amr3) or empty vector were agroinfiltrated (OD_600_=0.5) in *N. benthamiana* plants. The site of infiltration was marked in the leaves and two days after agroinfiltration, *Pseudomonas syringae* D36E carrying the respective effector was infiltrated in the marked area on the leaf. Cell death was observed 3 days post *P. syringae* pv. *tomato* DC3000, D36E infiltration. (F) Protein accumulation in transgenic Arabidopsis plants. Schematic representation of constructs transformed in Arabidopsis that contain either sensor and helper or sensor only and validation of its expression in T2 transgenic lines. Anti-FLAG and Anti-MYC immunoblot respectively shows accumulation of sensor and helper NLRs. Red arrow indicates protein band corresponding to the either sensor or helper NLR. Panel below shows Ponceau S staining of Rubisco as a loading control. *Indicates a non-specific band. (G-I) Effector recognition by Solanaceae sensor-helper pair results in cell death in *A. thaliana*. (G) Independent T2 lines were infiltrated with *P. syringae* D36E that delivers by T3SS either AvrBs2, empty vector or AvrRpt2. Effector recognition by the Bs2/NRC2 sensor-helper pair resulted in cell death (indicated by red arrows-HR); whereas no cell death was observed in the sensor alone T2 plants (indicated by green arrows -no HR). Images were taken at 60 hours post infiltration (hpi). Panel on the right shows images of the same leaves taken under UV light. Cell death can be observed in green in the infiltrated area; in the absence of cell death the infiltrated area is red. (H) Occurrence of cell death upon effector recognition in independent T2 lines of Col-0 that express Bs2+NRC2. (I) Cell death in leaves was quantified by measuring electrolyte leakage. Data were collected from 2 independent experiments with at least 3 technical replicates per transgenic line. Error bars represent standard error of the mean (SEM).

**Supplemental Figure S3.**
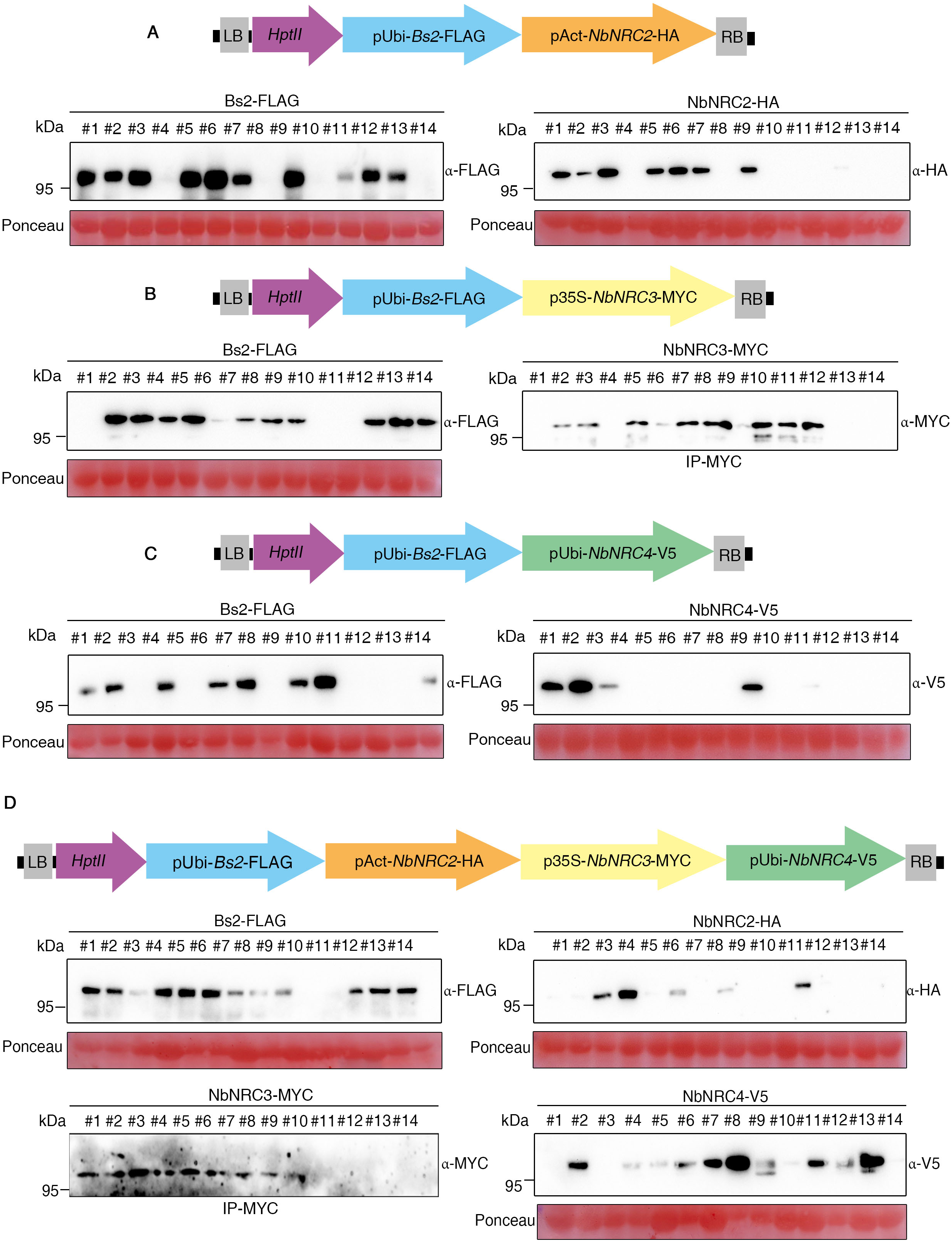
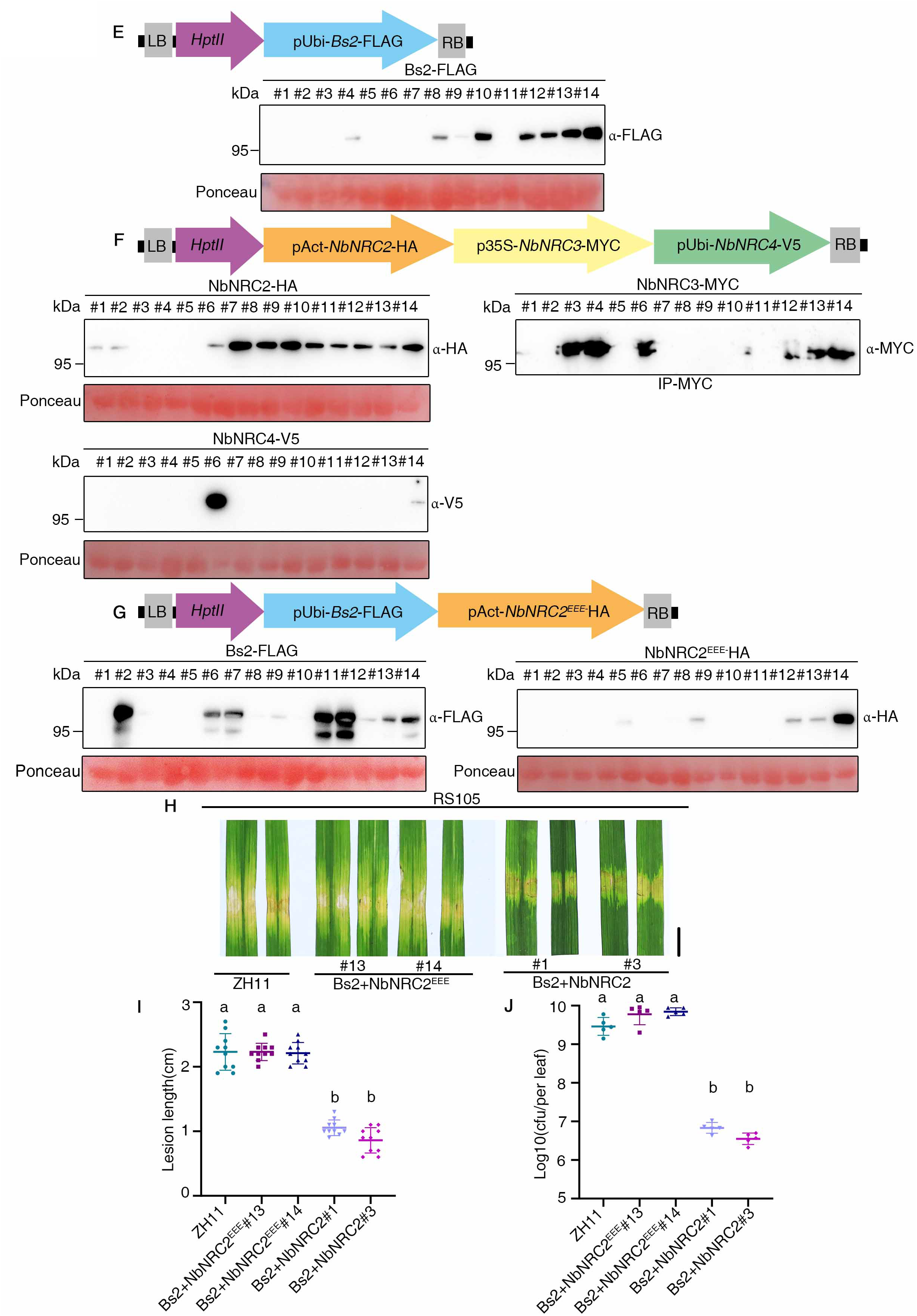
Bs2 confers resistance in transgenic rice against *Xoc* carrying AvrBs2 in a NRC2, NRC3, or NRC4 dependent manner. (A-C)Schematic representation of multi-transgene cassettes consisting of Bs2 and NRC helpers and validation of transgene expression in T1 transgenic plants. All individual transcription units were assembled into multigene constructs using the modular cloning system based on Golden Gate cloning. The arrows indicates the direction of transcription. A hygromycin phosphotransferase (*hptII*) selectable marker gene was used for transgenic plant selection. LB, left border; RB, right border. Bs2 with either NbNRC2 (A), NbNRC3 (B), or NbNRC4 (C). Immunoblotting below shows protein accumulation of Bs2 with either NbNRC2 (A), NbNRC3 (B), or NbNRC4 (C) in T1 transgenic rice generated in this study. Line names are given on the top. Total proteins were extracted from 2-week-old stable transgenic plants, separated in SDS–PAGE and immunodetected with the indicated antibodies. Ponceau S staining was used to demonstrate equal loading. (D) Schematic representation of construct containing Bs2 and a combination of the NRC-helpers NRC2/3/4. Validation of transgene expression of Bs2 and NRC-helpers NRC2/3/4 in T1 transgenic rice by immunoblotting below. (E-F)Schematic representation of construct containing only Bs2 or NRCs and validation of its expression in T1 transgenic lines. Schematic representation of construct containing Bs2 (E) and a combination of the NRCs (F). Immunoblotting below shows the protein accumulation of transgene expression of Bs2 and a combination of the NRCs in T1 transgenic rice. (G-J) An intact N-terminal MADA motif is required for NRC2-mediated resistance against bacterial leaf streak. Schematic representation of vector construct used for stable expression of Bs2 and MADA motif mutants of NRC2 (G).Validation of transgene expression of Bs2 and NRC2^EEE^ in T1 transgenic rice by immunoblotting(H). Comparisons of BLS disease resistance phenotype between Bs2/NRC2- and Bs2/NRC2^EEE^-coexpressed rice lines (I). Quantification of disease lesion length caused by *Xoc* in Bs2/NRC2- and Bs2/NRC2^EEE^-coexpressed rice lines (J). Lesion length was measured 14 days post-inoculation. Mean values and standard deviations were calculated from measurements of 10 independent leaves. Statistical significance is indicated by letters (p < 0.01, one-way ANOVA followed by Tukey’s post hoc test).

**Supplemental Figure S4.**
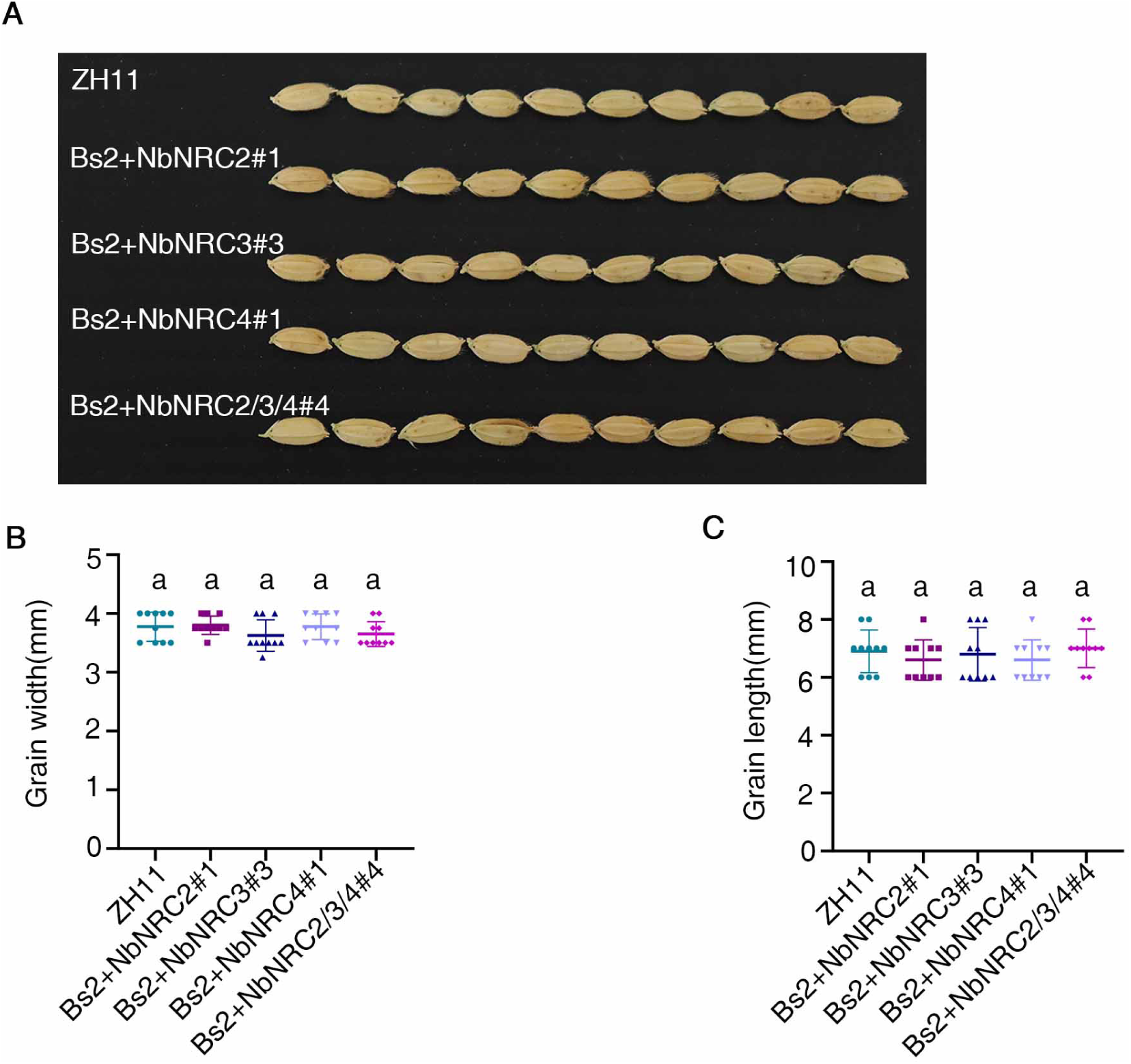
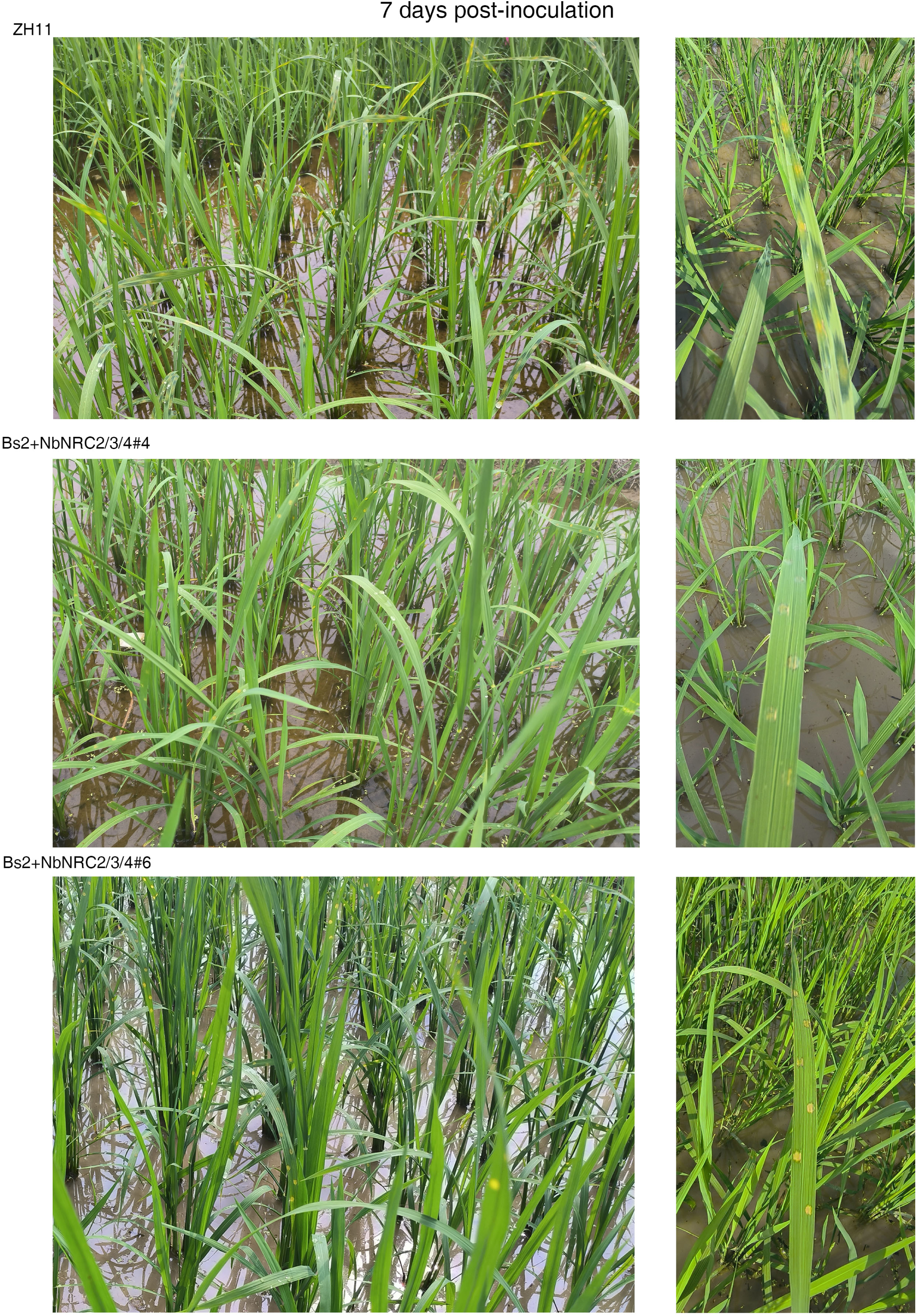
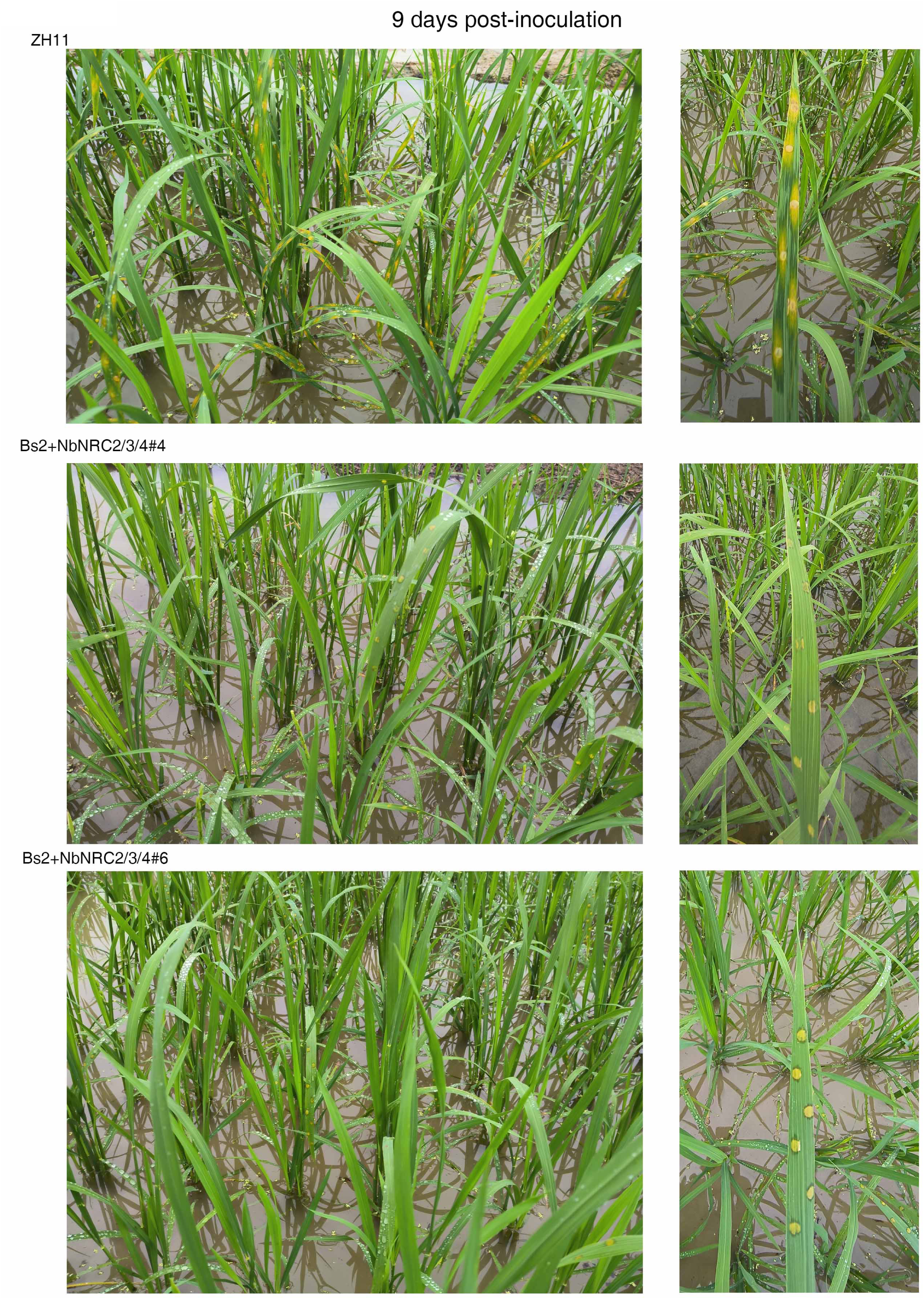
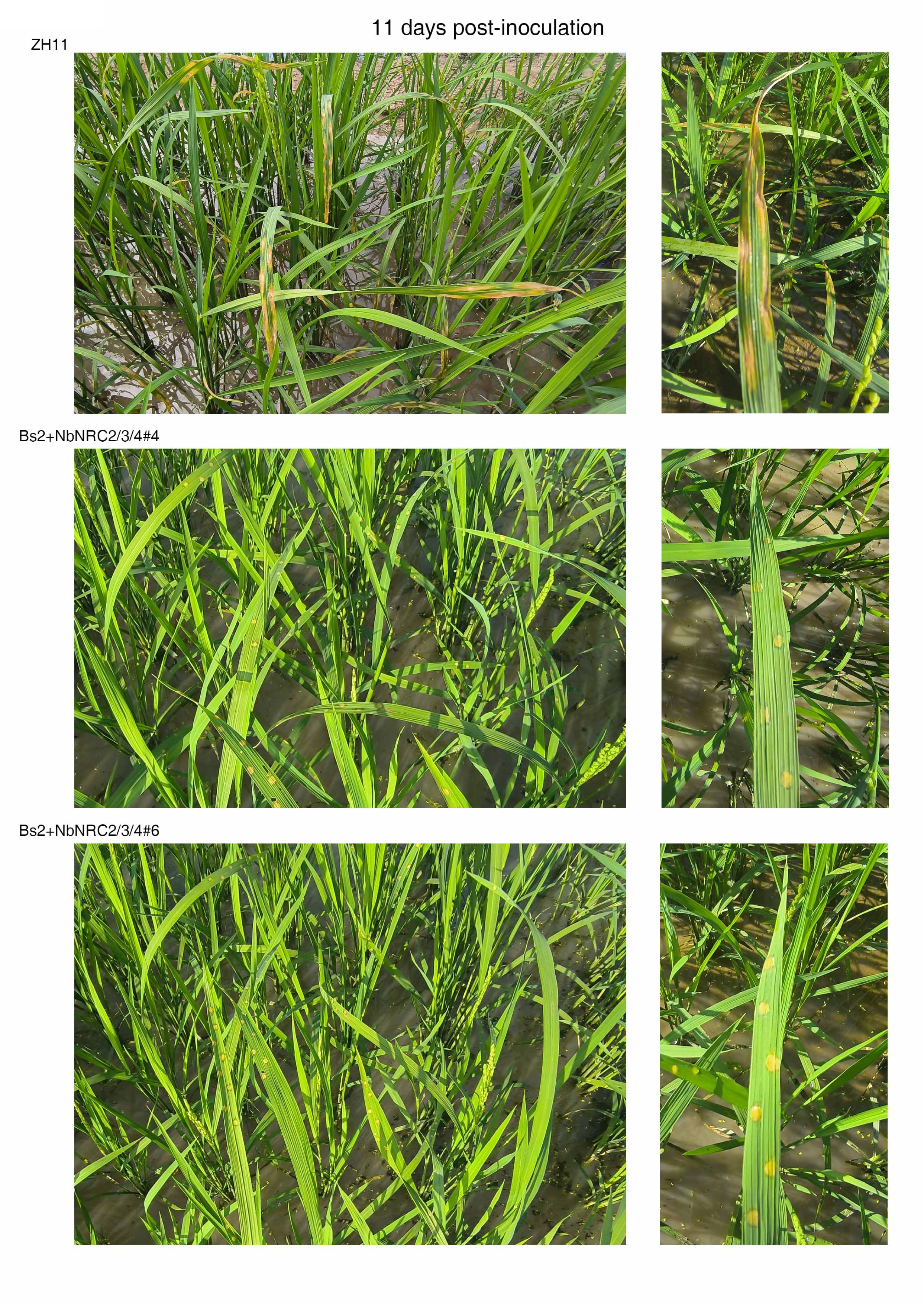
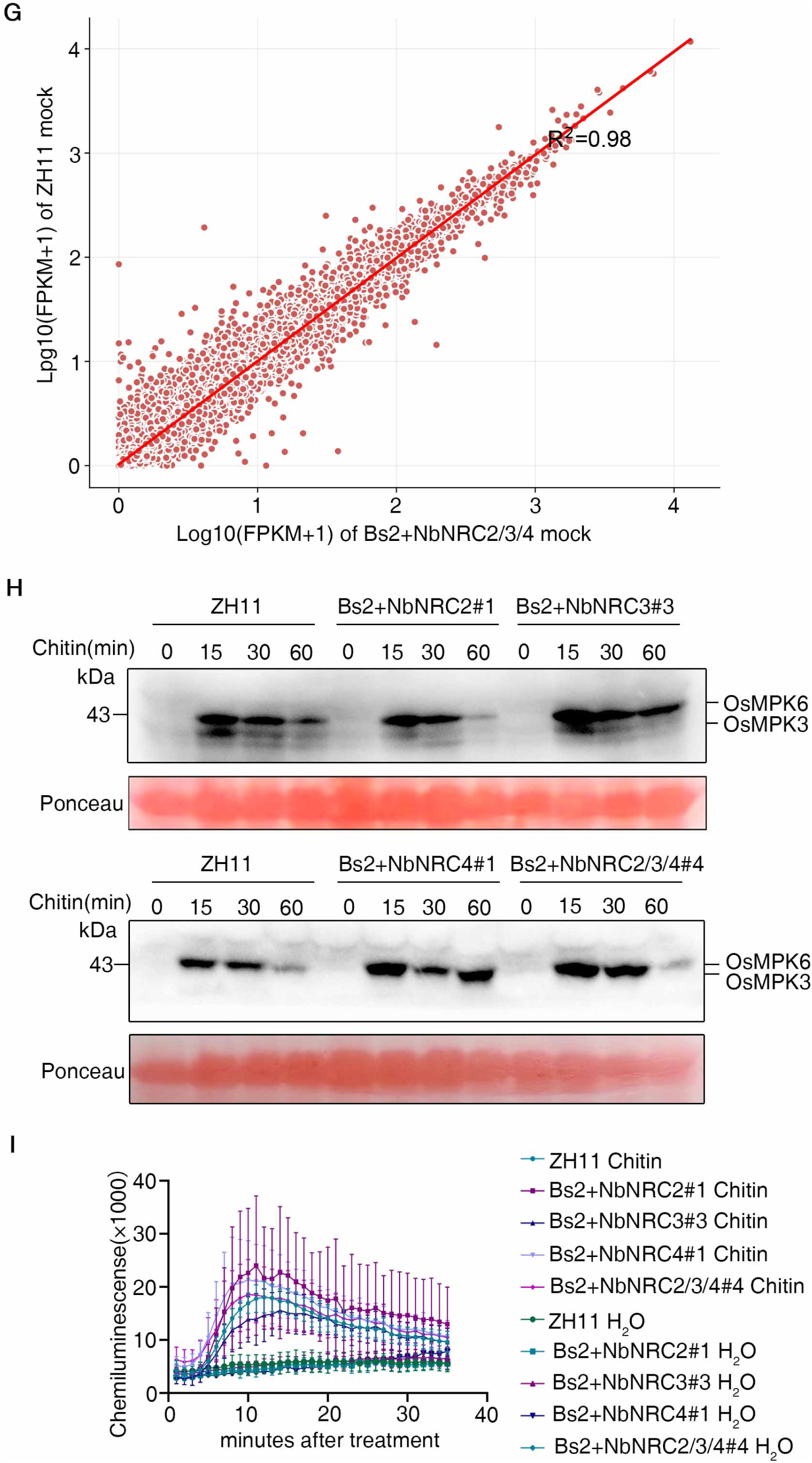
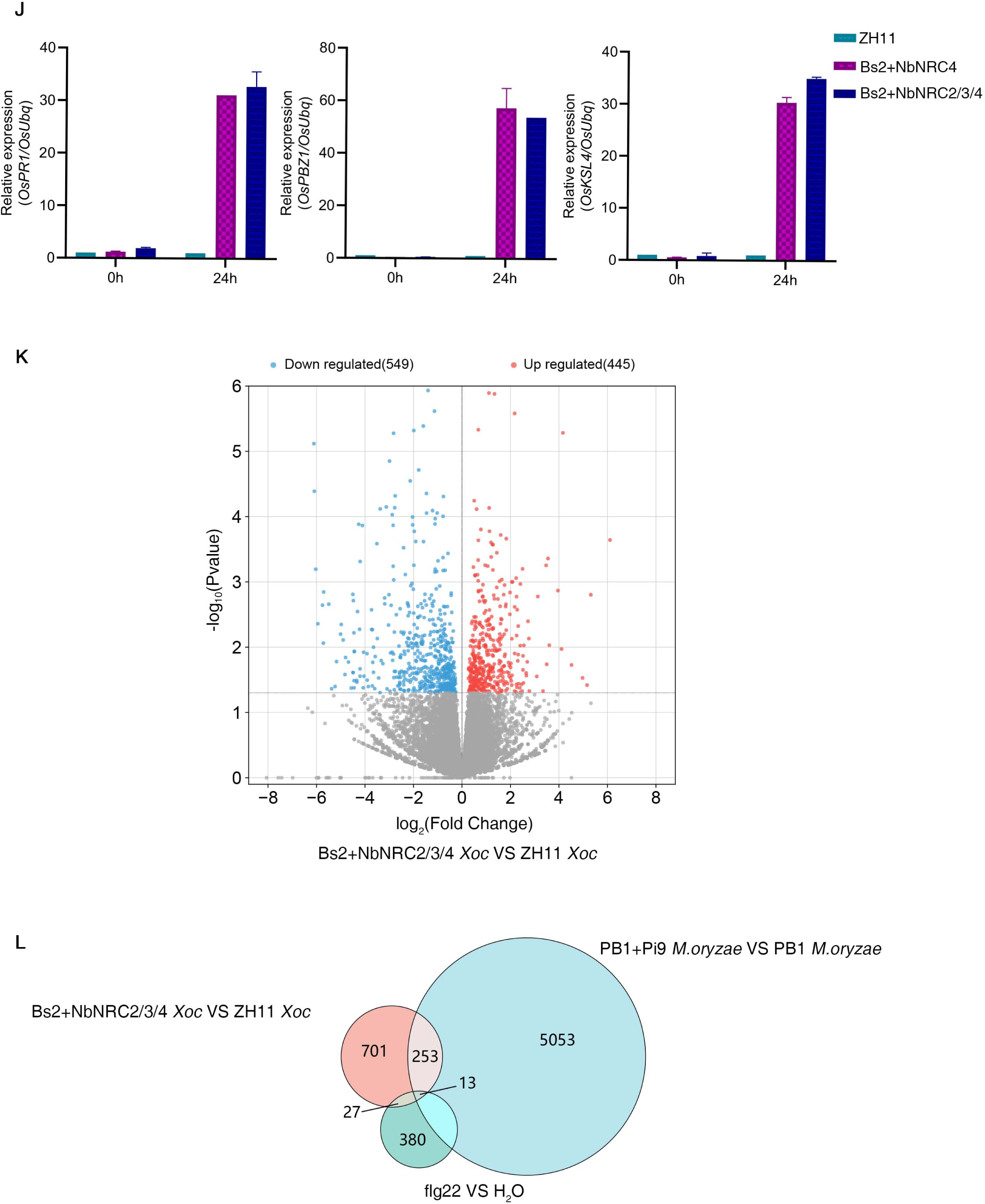
Heterologous expression of stacked senor and helper NLRs in rice does not affect basal resistance, growth and agronomic traits. (A-C) Comparisons of agronomic traits between wild-type rice ZH11 and different transgenic lines. Comparison of grain shape (A), grain width (B) and grain length (C) of ZH11 and different transgenic lines from the same planting conditions. (D-F) Bacterial leaf streak disease phenotypes of ZH11 and transgenic lines co-expressing Bs2 with triple combinations of NRC2/3/4 growing in the field. Comparison of disease phenotypes between ZH11 and transgenic lines planted in a standard field. Leaves of ZH11 but not transgenic plants showed characteristic water-soaking lesions expanding from the inoculation sites. Photos were taken at 7 dpi (D), 9 dpi (E), and 11 dpi (F). Individual leaves are enlarged and shown on the right side for each panel. (G) Co-expressing Bs2 with triple combinations of NRC2/3/4 do not significantly change transcriptome profiles of transgenic plants. (H) ROS generation induced by chitin treatment in transgenic lines coexpressing Bs2 with NRCs and ZH11 control. Leaf discs were treated with 10µg/ml chitin. ROS generation was monitored for 30 min. Values are means ± SD calculated from at least three technical repeats. (I) Chitin-induced MAPKs activation profile in the wild-type ZH11 and transgenic seedlings co-expressing Bs2 with NRCs. Ten-day-old seedlings were treated with 10µg/ml chitin and collected at the indicated time points. MAPK activation was analyzed by immunoblot analysis using phosphor-p44/42 MAPK (Erk1/2) (top panel), and the equal protein loading is indicated by Ponceau S staining for Rubisco (bottom panel). (J) Relative differential expression levels of the defense-related genes between ZH11 and Bs2+NRCs upon *Xoc* inoculation (K). The samples were collected at 0 hpi and 24hpi for qRT-PCR. Experiments were repeated three times and representative data are shown. (K-L)Transgenic rice co-expressing *Bs2* and *NRCs* displays NLR-induced transcriptome features upon *Xoc* infection. Volcano plot showing differentially expressed genes (DEGs) in Bs2+NRC2/3/4 compared with ZH11 infected with *Xoc*. A log2(fold change)>1 and q-value<0.005 were used as the cut off values (L). Venn diagram shows 266 and 40 overlapping genes between Bs2/NRCs-induced DEGs and Pi9-induced DEGs and flg22-induced DEGs, respectively (M). (p<0.05 and fold change>1)

## Notes

### Competing Interest Statement

The authors have declared no competing interest.

