## Supplementary Table S2 for "Interfamily co-transfer of sensor and helper NLRs extends immune receptor functionality between angiosperms"

**Table S2. Primers used for plasmid construction in this study.**

| Purpose | Gene/Fragments | Vector | Name | Sequence (5’-3’) |
| --- | --- | --- | --- | --- |
| Rice protoplast transient expression | AvrBs2*^Xoc^* | pRTV-nFLAG | AvrBs2^Xoc^-F | GATGATGATAAGGGATCCATGCGTATAGGTCCTCCGC |
|  |  |  | AvrBs2^Xoc^-R | ACTAGTAAGCTTGGTACCCTCCGGCTCGGTCTGGTTGG |
|  | Bs2 |  | Bs2-F | GATGATGATAAGGGATCCATGGCTCATGCAAGTGTG |
|  |  |  | Bs2-R | ACTAGTAAGCTTGGTACCATGTTCTTCTGAATCAGAATC |
|  | *Nb*NRC2 |  | *Nb*NRC2-F | GATGATGATAAGGGATCCATGGCGAACGTTGCGGTGG |
|  |  |  | *Nb*NRC2-R | ACTAGTAAGCTTGGTACCGAGATCGGGAGGGAATATAG |
|  | *Nb*NRC3 |  | *Nb*NRC3-F | GATGATGATAAGGGATCCATGGCAGATGTAGCAGC |
|  |  |  | *Nb*NRC3-R | ACTAGTAAGCTTGGTACCCAATCCGAGATCTGGAGG |
|  | *Nb*NRC4 |  | *Nb*NRC4-F | GATGATGATAAGGGATCCATGGCAGATGCAGTAGTG |
|  |  |  | *Nb*NRC4-R | ACTAGTAAGCTTGGTACCCTGTGTGGCCTTGGATCC |
| *N.benthamiana* transient expression | AvrBs2*^Xcv^* | pCAMBIA1300 | AvrBs2*^Xcv^*-F | CGGGGGACGAGCTCGGTACCATGCGTATCGGTCCTCTGCAAC |
|  |  |  | AvrBs2*^Xcv^*-R | ATCATGGTCTTTGTAGTCGTCGACATCCGTCTCCGTCTGCCTGG |
|  | AvrBs2*^Xoc^* |  | AvrBs2*^Xoc^*-F | CGGGGGACGAGCTCGGTACCATGCGTATAGGTCCTCCGC |
|  |  |  | AvrBs*^/Xoc^*-R | ATCATGGTCTTTGTAGTCGTCGACCTCCGGCTCGGTCTGGTTGG |
| qPCR | PR1a |  | PR1a-F | GTGTCGGAGAAGCAGTGGTA |
|  |  |  | PR1a-R | CGAGTAGTTGCAGGTGATGAAG |
|  | PBZ1 |  | PBZ1-F | AGGACTACCTCGTCGCTCAC |
|  |  |  | PBZ1-R | ACATTTCTGCGGCTCTCATT |
|  | KSL4 |  | KSL4-F | CGGTGTCATTCCTAAATCATGCAAGG |
|  |  |  | KSL4-R | CGGCCTGAGAGTAGAACACA |
|  | Ubi |  | Ubi-F | TTCTGGTCCTICCACTTTCAG |
|  |  |  | Ubi-R | ACGATTGATTTAACCAGTCCATGA |
|  | MoPot2 |  | MoPot2-F | ACGACCCGTCTTTACTTATTTGG |
|  |  |  | MoPot2-R | AAGTAGCGTTGGTTTTGTTGGAT |
